# Deep-Learning for Automated Markerless Tracking of Infants General Movements

**DOI:** 10.1101/2022.07.13.499984

**Authors:** H. Abbasi, S.R Mollet, S.A. Williams, L. Lim, M.R. Battin, T.F. Besier, A.J.C. McMorland

**Affiliations:** Auckland Bioengineering Institute (ABI), University of Auckland, Auckland, New Zealand, 1010, New Zealand; Liggins Institute, University of Auckland, Auckland, 1142, New Zealand; Department of Newborn Services, Auckland City Hospital, Auckland, 1142, New Zealand; Department of Exercise Sciences, Faculty of Science, University of Auckland, 1141, New Zealand

**Keywords:** neonatal general movement assessment (GMA), automated markerless motion tracking, deep-learning, DeepLabCut, convolutional neural networks (CNN), image processing, motion capture

## Abstract

The presence of abnormal infant General Movements (GMs) is a strong predictor of progressive neurodevelopmental disorders, including cerebral palsy (CP). Automation of the assessment will overcome scalability barriers that limit its delivery to at-risk individuals.

Here, we report a robust markerless pose-estimation scheme, based on advanced deep-learning technology, to track infant movements in consumer mobile device video recordings. Two deep neural network models, namely Efficientnet-b6 and resnet152, were trained on manually annotated data across twelve anatomical locations (3 per limb) in 12 videos from 6 full-term infants (mean age = 17.33 (SD 2.9) wks, 4 male, 2 female), using the DeepLabCut^™^ framework. *K*-fold cross-validation indicates the generalization capability of the deep-nets for GM tracking on out-of-domain data with an overall performance of 95.52% (SD 2.43) from the best performing model (Efficientnet-b6) across all infants (performance range: 84.32– 99.24% across all anatomical locations). The paper further introduces an automatic, unsupervised strategy for performance evaluation on extensive out-of-domain recordings through a fusion of likelihoods from a Kalman filter and the deep-net.

Findings indicate the possibility of establishing an automated GM tracking platform, as a suitable alternative to, or support for, the current observational protocols for early diagnosis of neurodevelopmental disorders in early infancy.

## 1. Introduction

### 1.1. Background

Injury to the developing fetal or infant brain (for example, due to hypoxic-ischemic encephalopathy, perinatal stroke, or infection) can cause severe impairments that lead to physical disability and cerebral palsy (CP) [1-3]. General Movements (GMs) are classes of movements observed in healthy infants up to around 5 months of corrected age [4, 5] where the quality of GMs reflects the health of the infants’ neuromotor system. Early prediction of neurological disorders facilitates early intervention during critical periods of heightened neuroplasticity, which a growing body of evidence confirms improved clinical outcomes [6].

The General Movements Assessment (GMA) is considered the most predictive motor assessment tool and a reference standard for determining the risk of CP [7-9] with a sensitivity of 98% [10]. Current protocols for manual GM scoring require the availability of at least two highly-trained clinicians, observing the infant (via video viewed at normal or increased speed) in an awake, calm, and alert state to evaluate the spatial and temporal complexity and variation of their movements [4, 11, 12]. Best prediction is achieved when the GMA is combined with MRI along with medical history for risk factors [4]. Nonetheless, the GMA is not yet widely adopted as a component of standard clinical care; one of the barriers to implementation is the need for trained assessors to perform GMA scoring. For those that are trained, manual assessment can be a time-consuming task, requiring breaks from assessors reviewing high numbers of videos to minimize fatigue-related error. To aid in streamlining finite clinical resources, often only infants meeting specific clinical criteria will undergo an assessment of their GMs, introducing the possibility of missed case identification of those not meeting the initial criteria. Hence, there is an urgent need to develop technological methods that can circumvent the bottleneck of the human assessor. Automated motion capture offers a low-cost, practical alternative to track and analyze anatomical movements effectively.

Automated neonatal GM tracking and analysis has been explored by different research teams using various techniques [13], including utilization of wearable biotech [14, 15], 3D motion capture [16, 17], wearable accelerometers [18], 3D RGB-D camera recordings [19], and conventional video recordings [20-25], and another deep-learning markerless pose estimation algorithm [30]. Commercial 3D motion capture systems (e.g., Vicon [26]) provide gold-standard spatial and temporal accuracies under laboratory settings but are impractically expensive and immobile. Wireless accelerometer sensors have also been used in a more controlled environment such as Neonatal Intensive Care Units (NICU) [27]. However, specialist equipment is again required and the attachment of physical devices to the infant may significantly impact their behavioral and physiological responses, as has already been shown with routine nursing interventions and even environmental noise [28]. To overcome the practical challenges in the studies above, simpler motion analysis strategies have been proposed to capture overall variations in spontaneous movements through assessing the motions of the body center of mass [29]. The latter approach does not, however, provide information about subtle movements such as torso rotations, which are important kinematic features considered in the GMA.

2D recordings from single-camera devices, such as web-cams, hand-held devices, and baby-monitors, provide lower spatial accuracies than gold-standard 3D motion capture systems. However, low resolution requirements for the capture of movement patterns, along with the broad availability of single-camera devices, reasonable price, and their flexibility of use (e.g., in home environments) make them an ideal, clinically relevant tool for capturing infant motor behavior. The utilization of technology readily available in the home creates the possibility of the assessment data collection being performed by parents, at the appropriate developmental stages, with little or no training, at convenient times, and with minimal disturbance of the infants’ natural physical and mental states. Robust automated analysis, including automatic movement tracking and infants GM classification, such as that proposed here, is required to process the extensive amount of produced data.

Deep neural network architectures are potent tools for reliably automating challenging signal processing tasks such as classification, identification, and segmentation. However, to achieve their potential, these structures require large datasets for authentic learning/training and robust performance. Dataset construction can be even more challenging when it relies on manual annotation of data completed by various observers, with associated biases [31]. The application of deep-learning-based approaches is relatively new in the infant GMA field, and only a few recent attempts have aimed to apply these novel techniques for movement classification, not motion detection in 2D videos [32-34].

This work, for the first time, proposes a robust deep neural network, developed in the DeepLabCut^™^ environment [35, 36] for automatic markerless motion tracking of infant movements, in standard iPad video recordings, for the purpose of detecting GMs in the fidgety period. The approach’s generalization performance will be assessed using k-fold cross-validation across 12 video recordings from six infants.

Ease of data collection will facilitate the collection of clinically useful databases of kinematic recordings which, coupled with on-going records of neurological outcomes, will be essential for developing predictive machine-learning models that could robustly assist with early risk assessments. Harnessing the recent enhancements in image processing could help to establish automated platforms that can accurately analyze GM kinematics and monitor neonates’ neurological development continuously [37].

## 2. Data acquisition

### 2.1. Ethics

All procedures in this study were approved by the Auckland Health Research Ethics Committee (AHREC 000146). Parents/caregivers were fully informed of the experimental purpose, filming procedure and methods, and provided informed written consent for their child’s participation.

### 2.2. Clinical procedures

Infants were recruited using social media, flyers and posters. A cohort of six infants (four males and two females, mean height 62.66 (SD 2.56) cm, weight 6.98 (SD 0.24) kg) were assessed within the fidgety period: mean age of 121.3 (SD 20.4) days post-term corrected age (equal to 17.33 (SD 2.92) weeks) (Table 1). After family had provided written consent, infants were brought to the lab, changed into a nappy only, and laid down in a supine position on a white cotton mat. Two standard mini-iPads (1^st^ device: iPad mini 5^th^ generation, software version:14.4.2, model No. MUQX2X/A, camera resolution: 8-megapixel and 2^nd^ device: iPad mini 4, software version:13.4.1, model No. MNY32X/A, camera resolution: 8-megapixel 1080p HD video recording) were set up on tripods on both left and right sides of the infant. Infants were filmed while awake with spontaneous mobility in their natural state for 3–5 minutes with no external distractions such as from their parents/caregivers or the presence of toys.

**Table 1.** Infants’ health information

|  | **P01** | **P02** | **P03** | **P04** | **P05** | **P06** | **Mean (SD)** |
| --- | --- | --- | --- | --- | --- | --- | --- |
| **Age (days)** | 141 | 104 | 87 | 124 | 146 | 126 | 121.3 (20.4) |
| **Sex** | F | M | M | M | F | M |  |
| **Ethnicity** | NZE*/<br>Māori | NZE/<br>Chinese | NZE | NZE | NZE/<br>Cook Island Māori | NZE/<br>Chinese |  |
| **Height \*\*** | 62 | 60 | 59 | 66 | 65 | 64 | 62.7 (2.6) |
| **Mass (kg)** | 6.8 | 7.5 | 6.9 | 6.9 | 7.0 | 6.8 | 7.0 (0.2) |
| **Torso (Length)** | 22 | 10 | 21 | 21 | 26 | 22 | 20.3 (4.9) |
| **Torso (Width)** | 18 | 7 | 16 | 15 | 14 | 17 | 14.5 (3.6) |
| **Upper arm** | 8 | 10 | 9 | 12 | 11 | 10 | 10.0 (1.3) |
| **Forearm** | 7 | 7 | 9 | 9 | 10 | 9 | 8.5 (1.1) |
| **Upper leg** | 12 | 12 | 11 | 13 | 13 | 12 | 12.2 (0.7) |
| **Lower leg** | 11 | 10 | 11 | 13 | 13 | 12 | 11.7 (1.1) |
| **Head** | 14 | 18 | 14 | 17 | 14 | 13 | 15.0 (1.8) |
| **Right Hand** | 5.5 | 7 | 7 | 7 | 8 | 8 | 7.1 (0.8) |
| **Right Foot** | 9 | 8 | 8 | 8 | 8 | 8 | 8.2 (0.4) |

\* New Zealand European
\*\* All length measurements are given to the nearest 0.5 cm, and presented in cm.

A total of 12 videos (resolution: 1920×1080 pixels, frequency: 29.97 frames/second) with an overall number of more than 69k frames were captured from the six infants. Videos were post-processed in Adobe Premiere Pro 2021 by situating the infant in the center of the frame and cropping the remaining excessive part of the image out. Videos were then saved in their original quality. These preprocessing steps removed extra unnecessary information in the videos, which was anticipated to improve the robustness of the model. Twelve anatomical locations (3 per limb: *shoulders, elbows, hands, hips, knees*, and *feet)* were manually labeled by an expert (HA) across 2400 frames in the entire training set (equal to 28,800 points). The 2400 frames included 200 frames from each video that were automatically extracted from each recording using k-means clustering implemented within DeepLabCut to maximize frame diversity. Examples of the manually labeled locations are shown with colorful dots in Fig. 1.

**Figure 1.**
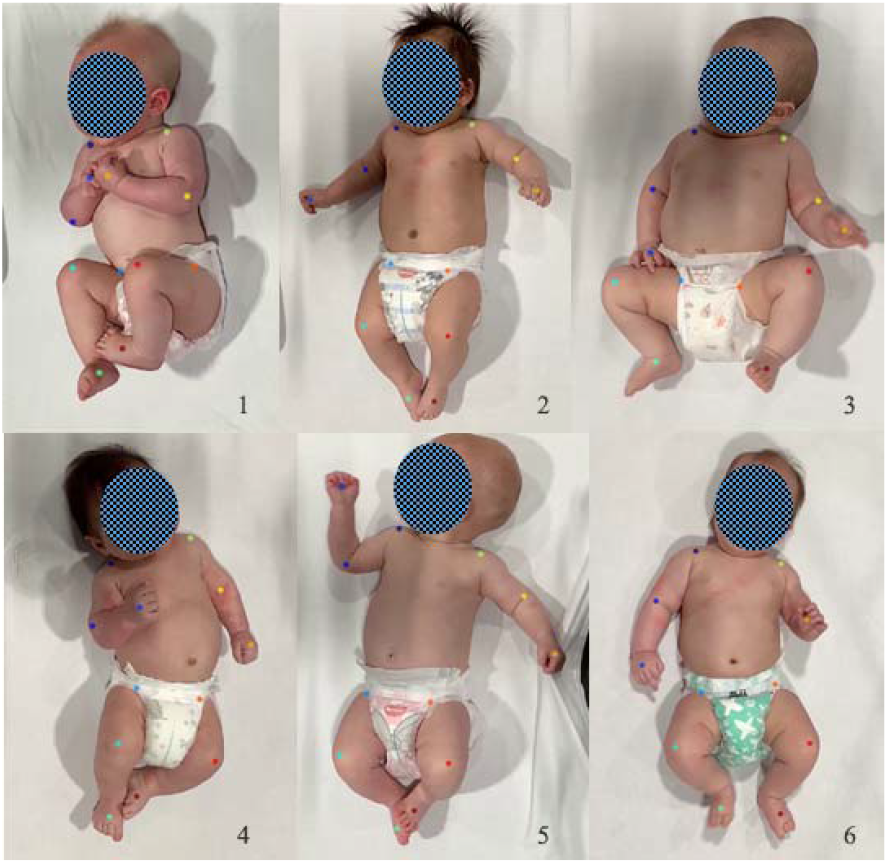
Examples of the twelve manually labeled anatomical locations (colorful dots) in six different infants in DeepLabCut environment.

## 3. Methods and computational approach

### 3.1. Deep-Learning approach

Enhancements in computer science and advanced image processing, including deep-learning, have allowed researchers to develop multiple open-source motion-tracking toolboxes, such as DeepLabCut^™^ and OpenPose [38], that can follow specified points across input frames. DeepLabCut [36] is a deep-learning-based platform for markerless pose estimation with successful performance in the movement tracking of various species, including non-primates such as cheetahs, horses [39], and mice as well as primates such as macaques [40]. Nath et al. have also reported the possibility of using this platform for human infant motion tracking [35, 41]. Recent reports indicate the robust generalization capability of DeepLabCut in movement tracking of out-of-domain animal subjects (horses) with normalized errors close to the within-domain tested subjects [39].

DeepLabCut has been reported to achieve its excellent performance by combining key elements of deep convolutional networks through using 1) deep resnets and efficientnets that are pretrained on a benchmark ImageNet [42] used for object recognition, and 2) deconvolutional layers, instead of a classification layer, for semantic segmentation [36]. Resnet-152 is a top-performing deep-net model created by Microsoft for the ImageNet challenge in 2016 [43]. These types of deep artificial neural networks are generally designed to mimic the mechanism of certain biological neuronal cells, pyramidal neurons, to improve learning by optimally skipping intermediary connections in a deep structure [44]. On the other hand, transfer learning in efficientnet architecture classes [39] uses a compound scaling method through uniform scaling of the network’s depth, width, and resolution. This strategy has been shown to out-perform the resnet models on out-of-domain ImageNet data [39]. A recent study suggests that the efficientnet-b6 can achieve better accuracies than other efficientnet classes (i.e., b0 to b5) [39]; hence we used this model in the current study. For simplicity, we will call the resnet-152 and efficientnet-b6 models ‘resnet’ and ‘efficientnet’ from here on. Deconvolutional layers are known to up-sample the visual details and generate probability densities in spatial space which can be later used as evidence to locate a specific body part in a particular location. In DeepLabCut, the deep-nets are iteratively fine-tuned by updating/adjusting the weight parameters using the manually labeled data. DeepLabCut automatically assigns high probabilities to the labeled anatomical sites and allocates low-likelihoods to the rest of the image [36].

In this work, we first utilize the DeepLabCut environment to develop robust deep-learning models for markerless motion tracking in human infants’ video recordings (resnet training: DLC version 2.1.10.2, TensorFlow: 1.14, CUDA: 11.3, Python 3.7.9; efficientnet training: DLC version 2.2.0.2, TensorFlow: 2.5.0, CUDA: 11.4, Python 3.8.6; Efficientnets became available in the later version of DLC). Furthermore, this work aims to assess how well the model can generalize across all infants through a leave-one-out *k*-fold cross-validation strategy across data from all infants.

### 3.2. *k*-fold cross-validation

A subject-based *k*-fold cross-validation strategy [45] can implicitly specify whether a model performs equally well across all subjects and helps to identify whether there is a significant variation in the dataset. In other words, training of the networks over *k -* 1 (five, in this case) infants and testing that on the data from an unseen infant can determine the degree of reliability of the proposed motion tracking scheme. Here, we considered a 6-fold subject-based cross-validation strategy for the six infants in the dataset to assess how well the model generalizes across all infants with various movements. The terms ‘testing’ and ‘validation’ are used somewhat interchangeably in the literature, which can cause confusion. In the following discussion we follow the conventions used by DeepLabCut, using the term ‘testing’ to refer to data withheld from the training to assess performance and update training hyperparameters during learning process, whereas the word ‘validation’ is used to refer to a dataset that is withheld to the very end of the process for performance evaluation. In each training fold, data from one particular baby were left out, and the two recordings associated with that baby were excluded from the training set. This strategy allows comparison and validation of the learning schedule for, and between, the resnet-152 and efficientnet-b6 models.

A 95%-to-5% training/testing scheme was used to train the deep-nets using an *imgaug* image augmentation strategy [46] in each training fold (round). The learning rate was initially set at 0.005 and decreased to 0.001 through a recommended multi-step updating regime of [0.005, 0.02, 0.002, 0.001] for up to [10,000, 430,000, 730,000, and 1,030,000 iterations], respectively. The convergence of the cross-entropy loss function during training was confirmed across all 6-folds with a sharp decay during the first 30,000 iterations (for both models) and then gradually decayed further to lower values during 1,030,000 and 700,000 iterations for the resnet and efficientnet, respectively. As mentioned earlier, efficientnet is a larger network where 700k training iterations were performed, taking over 7 days, as compared to the 1,030k iterations for the resnet. The root mean square errors (RMSE), namely the average distance between the identified labels by the resnet classifier (DLC) and the scorer’s identifications, were evaluated at each snapshot throughout the training process across all six-folds, resulting in an overall training error of 2.73 (SD 0.06) pixels at the 1,030k iteration for the resnet and 7.77 (SD 0.11) pixels at the 700k iteration for the efficientnet. Training took 126 (SD 3.5) hours across the six-folds on the high-performance machines detailed in the “computing infrastructure” section. In contrast, test errors were found to be higher at 6.73 (SD 0.17) pixels and 9.59 (SD 0.32) pixels for the resnet and efficientnet, respectively. These findings are consistent with previously described capabilities of DeepLabCut [36]. Fig. 2 demonstrates evaluated RMSEs (pixel) across all six folds for training over 1,030k and 700k iterations for the resnet and efficientnet. This plot confirms the fast fine-tuning of the models across all folds. After each training fold, the performance of the model was evaluated (validated) on the two videos from the excluded infant (out-of-domain/unseen participant). This cross-validation procedure was repeated six times across data from six babies, each time removing one infant’s data from the training set and using that excluded infant’s data for testing the resulting trained network.

**Figure 2.**
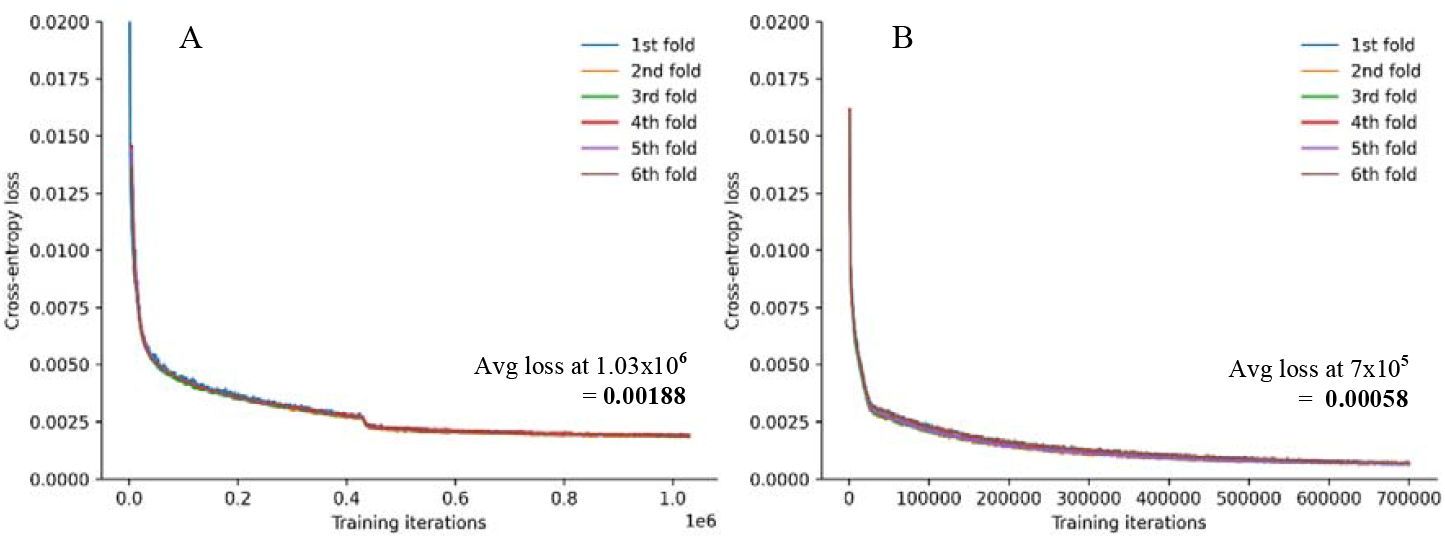
Evaluated cross-entropy loss for all training folds using (A) resnet-152 and (B) efficientnet-b6.

### 3.3. Kalman filter for an aPCK-based performance measure

To quantify the performance of markerless pose-estimation, automatically determined marker positions are needed to be compared against labels determined in another way, usually from manual labelling. To compare between evaluation metrics, the literature has established a trustable criterion, called the average Percentage of Correct Keypoints (aPCK) [39, 47, 48]. The aPCK approach needs a human assessor to manually annotate a validation set (on the top of training/test set). The aPCK then labels a predicted marker as ‘correctly detected’ if its location falls within a certain circular distance to the ground-truth manually annotated location. The distance is usually chosen as a fraction of an image-specific scaling factor such as the torso length. Because manual labelling of frames requires time-consuming, high precision work, this approach is challenging when the validation set includes large datasets and numbers of frames. One mitigation of this problem is to validate against only a small number of additionally labelled frames. Here, we present an alternative novel automatic approach that allows for evaluation against an entire video. In our case, this would be approximately equivalent to manual labelling ∼800k datapoints across ∼60k frames. Conceptually, our proposed strategy follows a similar approach to aPCK, but instead of requiring manual measures at each frame it uses the Kalman filter framework, building on the expectation of smooth and continuous motion, to estimate the position of a marker at time *t* based on the state (position, velocity, and acceleration) of the point in the previous frame *t* - 1. The Kalman filter uses automatically determined marker positions in previous frames as noisy measurements to update joint probability distributions for its position, velocity, and acceleration state variables, from which to predict a position in the current frame (Eqn 1) [49].

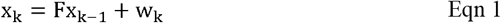

where the state vector *x*_*k*_ = [*d*_*X*_ *d*_*y*_ *v*_*X*_ *v*_*y*_ *a*_*X*_ *a*_*y*_]^T^ comprising x and y displacement, velocity and acceleration, the transition matrix 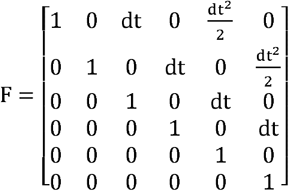, w_k is a noise term, and *dt* is the time interval between frames. The predicted marker positions can be used in place of the ground truth manually annotated locations traditionally used in aPCK for estimation of the motion tracking performance.

After each prediction step, the prediction is updated with a measurement of the current observed variables, the current marker position determined by the deep-net models from DeepLabCut, weighted by the calculated certainty of the current state and observation. The Kalman filter also returns the likelihood of each measurement, depending on its relationship to the probability distribution of the expected state. On the other hand, the classifiers from DeepLabCut (DLC), namely the resnet and efficientnet models, return a probability (p) that indicates their confidence level for each of the predicted anatomical locations (markers) at the identified location on the image (DLC-p). Here we propose to combine likelihoods from the Kalman filter with the predictions from deep-nets to identify true positive (TP), false positive (FP), false negative (FN), and true negative (TN) detections and automatically evaluate performance metrics for the markerless approach. The details of this approach are explained below. Through manual visual inspection and testing, a negative loglikelihood (KF-LL) greater than 20 was determined as a threshold above which observations were considered poor tracking. Similarly, we define good labelling when the DLC classifier has identified a marker with a confidence level of >= 0.6.

In our confusion matrix, logical combinations of log-likelihoods from Kalman filter represent ground-truth (equivalent to manual annotations in aPCK method) and predictions from the deep-net classifier (i.e. the resnet or efficientnet) serve as predicted measures. Observations from the forward Kalman Filter with likelihoods above a certain threshold [50] are not alone suitable to define ground-truth because the Kalman filter log-likelihood at each data step is influenced by the state(s) from the previous data steps, derived from incorrect DLC tracking. Additional information about the correctness of labelled locations can be derived by assuming that large contiguous blocks of low KF-LL values are correct, and that points that are neighbors to correct values, and with low KF-LL values from running the Kalman filter either forwards or backwards through the data, are also correct.

Figure 3 and Table 2 present a segment of data to illustrate how this approach handles DLC errors in two scenarios. In scenario 1 (black series in Fig. 3B), data point 2 is the only tracking error. However, datapoint 3 and 4 also have KF-LL > 20 when run in the forward direction despite being correctly tracked. Point 2 is correctly identified as the only incorrectly tracked point where both the forward and backward KF-LLs exceed the error threshold. In scenario 2 (orange series in Fig. 3B), datapoints 2 to 4 are incorrectly tracked and datapoint 3 is missed in the forward Kalman filter evaluations, due to its proximity to point 2. In scenario 2, datapoints < 1 and > 6 are labelled good because they are in large blocks where both the forwards and backwards KF-LL values are < 20. Points 1, 5, and 6 have only one KF-LL value > 20 and neighbor a good data point, so are also labelled as correct. Points 2, 3, and 4 remain labelled as poor tracking.

**Figure 3.**
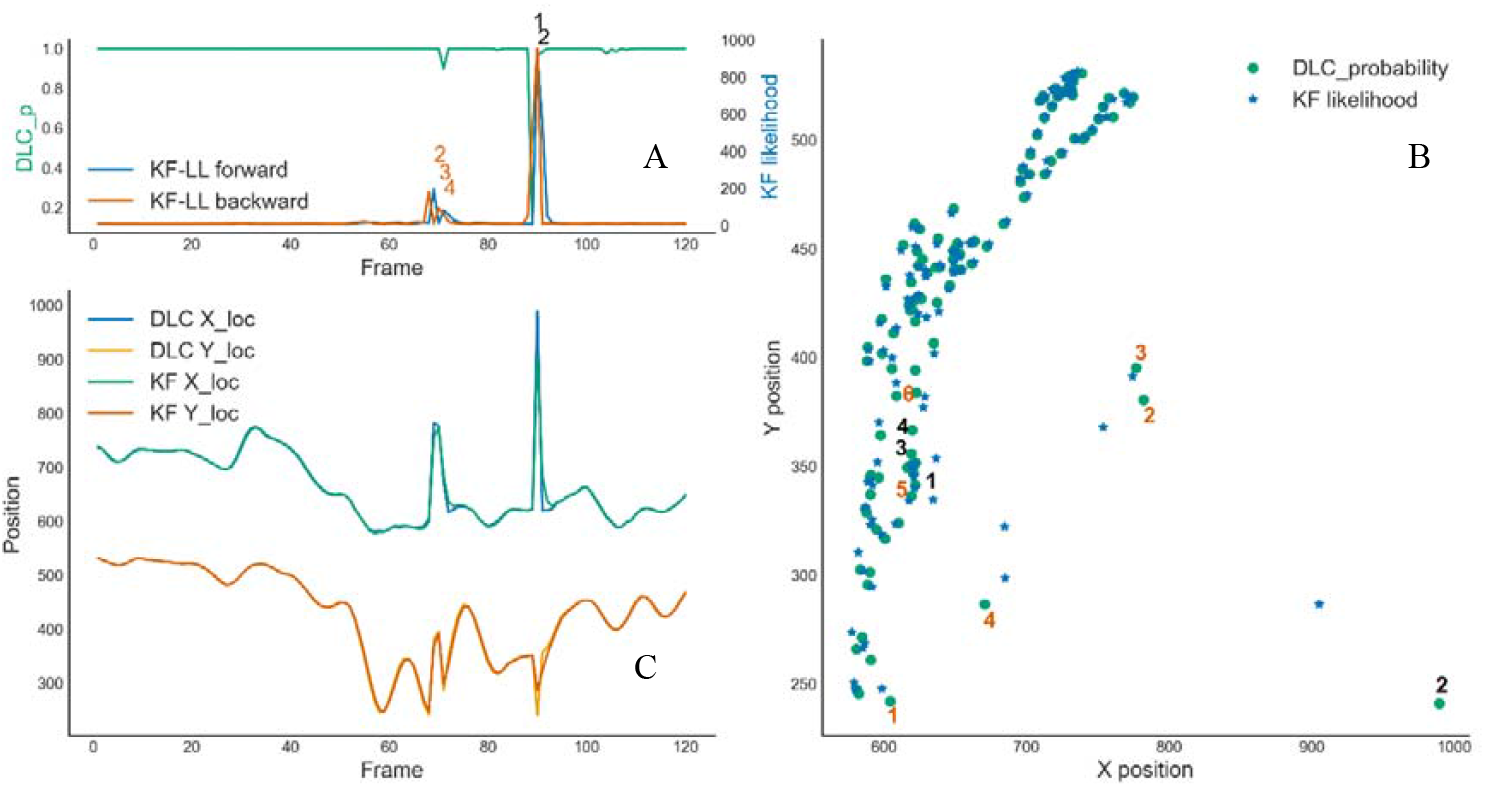
Example tracking errors A: Outputs of DLC-p (green), forward KF_LL (blue), and backward KF_LL (orange). B: 2D movement trajectories (scenario 1: black numbers, scenario 2: orange numbers). C: time-series of movement trajectories.

**Table 2.** Examples of DLC, forward Kalman filter, and backward Kalman filter outputs in two different tracking error senarios.

|  | Data point | DLC measures |  |  | Forward Kalman filter |  |  | Backward Kalman filter |  |  | Tracking | State |
| --- | --- | --- | --- | --- | --- | --- | --- | --- | --- | --- | --- | --- |
|  |  | X (px) | Y | DLC-p | X | Y | KF-LL | X | Y | KF-LL |  |  |
| Scenario 1<br>(black numbers) |  | 620.2 | 349.7 | 1.00 | 620.7 | 348.6 | 6.7 | 624.1 | 346.4 | 8.9 | Good | TP |
|  |  | 619.4 | 350.9 | 1.00 | 619.7 | 350.1 | 6.5 | 635.6 | 339.9 | 43.9 | Good | TP |
|  | 1 | 622.3 | 351.8 | 0.12 | 621.8 | 351.4 | 6.5 | 686.3 | 319.5 | 527.2 | Good | FN |
|  | 2 | 989.5 | 240.9 | 0.96 | 905.4 | 286.6 | 922.2 | 904.1 | 288.1 | 954.6 | Poor | FP |
|  | 3 | 619.5 | 355.9 | 0.98 | 684.7 | 322.3 | 550.0 | 618.1 | 358.3 | 7.3 | Good | TP |
|  | 4 | 619.9 | 366.9 | 1.00 | 636.7 | 353.7 | 50.2 | 618.8 | 370.3 | 8.2 | Good | TP |
|  |  | 622.7 | 383.9 | 1.00 | 627.7 | 377.1 | 13.9 | 622.9 | 387.8 | 9.2 | Good | TP |
|  |  | 634.9 | 406.8 | 1.00 | 635.5 | 401.7 | 10.7 | 634.0 | 410.1 | 8.0 | Good | TP |
| Scenario 2<br>(red numbers) |  | 591.0 | 260.9 | 1.00 | 586.7 | 268.5 | 14.9 | 599.6 | 256.6 | 15.9 | Good | TP |
|  | 1 | 604.6 | 242.0 | 1.00 | 598.8 | 247.6 | 12.7 | 630.4 | 255.8 | 179.8 | Good | TP |
|  | 2 | 782.3 | 380.6 | 1.00 | 753.9 | 368.0 | 193.0 | 777.2 | 382.9 | 9.6 | Poor | FP |
|  | 3 | 776.8 | 395.2 | 1.00 | 774.4 | 391.3 | 11.0 | 758.1 | 386.2 | 91.1 | Poor | FP |
|  | 4 | 670.8 | 286.7 | 0.90 | 685.0 | 298.5 | 80.2 | 653.5 | 303.7 | 63.7 | Poor | FP |
|  | 5 | 616.6 | 349.5 | 1.00 | 634.6 | 334.7 | 58.6 | 613.3 | 358.6 | 18.8 | Good | TP |
|  | 6 | 621.9 | 394.3 | 1.00 | 628.8 | 381.9 | 28.9 | 619.7 | 400.7 | 12.6 | Good | TP |
|  |  | 626.3 | 427.2 | 1.00 | 629.9 | 418.4 | 17.6 | 624.9 | 430.3 | 7.9 | Good | TP |
|  |  | 627.1 | 445.4 | 1.00 | 629.4 | 440.1 | 10.4 | 626.8 | 444.2 | 6.7 | Good | TP |

A state-machine approach to implementing the logic presented above is indicated below in pseudocode:

Read data: columns of DLC probability (DLC-p), forward KF (KF-LL_f_), backward KF

(KF-LL_b_)

Set thresholds values: KF-LL threshold (KF-T = 20), DLC-p threshold (DLC-T = 0.6),

…

number of points to assume is a good block (N = 10)

Label all points as *Poor* tracking

Mark as *Good* points in all blocks of >= N points long, where KF-LL_f_ < KF-T and KF-

LL_b_ < KF-T

Until no more changes are made, repeat over the following:

Amongst remaining *Poor* points, mark as *Good* all points i where…

KF_LL_f_[i] < KF-T and point i-1 is *Good*.

Amongst remaining *Poor* points, mark as *Good* all points i where…

KF_LL_b_[i] < KF-T and point i+1 is *Good*.

Mark point as TP, if DLC-p > DLC-T and point is *Good*

Mark point as FP, if DLC-p > DLC-T and point is *Poor*

Mark point as FN, if DLC-p < DLC-T and point is *Good*

Mark point as TN, if DLC-p < DLC-T and point is *Poor*

In our experience, a common situation where a FN can arise is when a body-part, in particular hand and foot, is either entirely or partially hidden behind or under other body-parts. In this case, the deep-net has correctly identified the body-part location (the good tracking criterion is met), but with sufficient uncertainty to classify the body-part as “unidentified” (DLC-p < 0.6).

This automatic performance evaluation approach allows the validation of larger markerless datasets. Here, we report the overall performance (Eqn 6) as the average of precision (positive predictive value (PPV) – Eqn 4) and sensitivity (true positive rate (TPR) – Eqn 5) in each fold or at each anatomical location. We will also evaluate accuracy (ACC – Eqn 7) to simultaneously consider FPs and FNs to identify which body-parts are associated with the poorest identification outcomes across all anatomical locations.

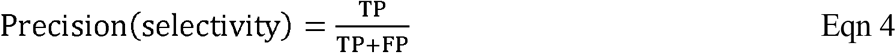

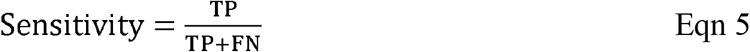

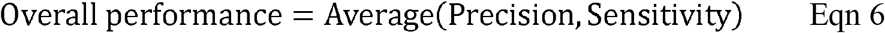

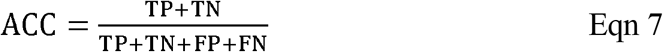

### 3.4. Computing infrastructure

The deep-net models, namely resnet-152 and efficientnet-b6, were trained using New Zealand eScience Infrastructure (NeSI) high-performance computing facilities’ Cray CS400 cluster [51]. The training process was executed using enhanced NVIDIA Tesla P100 GPUs with 12 GB CoWoS HBM2 stacked memory bandwidth at 549 GB/s (for resnet on DLC version 2.1.10.2) and NVIDIA Tesla A100 PCIe GPUs with 40 GB HBM2 stacked memory bandwidth at 1555 GB/s (for efficientnet on DLC version 2.2.0.2), per training task. The larger size of the efficientnet model and the associated necessary memory size, required us to utilize A100 GPUs instead of the P100s. Intel Xeon Broadwell CPUs (E5-2695v4, 2.1 GHz) were used on the cluster for handling the GPU jobs. The algorithms were run under Python environments (Python 3.7.9, pykalman 0.9.5).

## 4. Results

### 4.1. Fold-based vs location-based performance metric

The 6-fold cross-validation of the deep-net classifiers (trained on ten videos from 5 infants, tested on two videos from a novel infant) demonstrated a successful overall performance of 95.52% (SD 2.43) for marker tracking across all out-of-domain infants (subject-based metrics) using the efficientnet model, while the resnet resulted in an overall performance of 94.94% (SD 2.63). To help with a better understanding of the performance with unseen data, we also calculated performance for each anatomical location in the validation sets (Fig. 4). The heatmaps in Figures 4A-B and 4C-D show the precision and sensitivity measures associated with each anatomical location for each fold from the efficientnet and resnet models, respectively. The reported overall performance metrics in this article are the average of the values (mean %, SD) calculated from twelve evaluations. Figure 5 demonstrates how the accuracy of resnet and efficientnet models vary across anatomical locations. As these measures include the simultaneous impact of FPs and FNs, they help to specify which anatomical locations were more challenging for a certain deep-net classifier to identify and where exactly the model performed better. Results in Fig. 5 highlight that performance was poorest for tracking of hands and hips, while both models achieved their best tracking results for the left shoulder, right knee, and right foot. Tables S1 and S2 provide performance details for resnet and efficientnet, respectively. Figure S1 further provides a detailed schematic that compares the performance of the resnet vs efficientnet models in the subject-based and location-based schemes.

**Figure 4.**
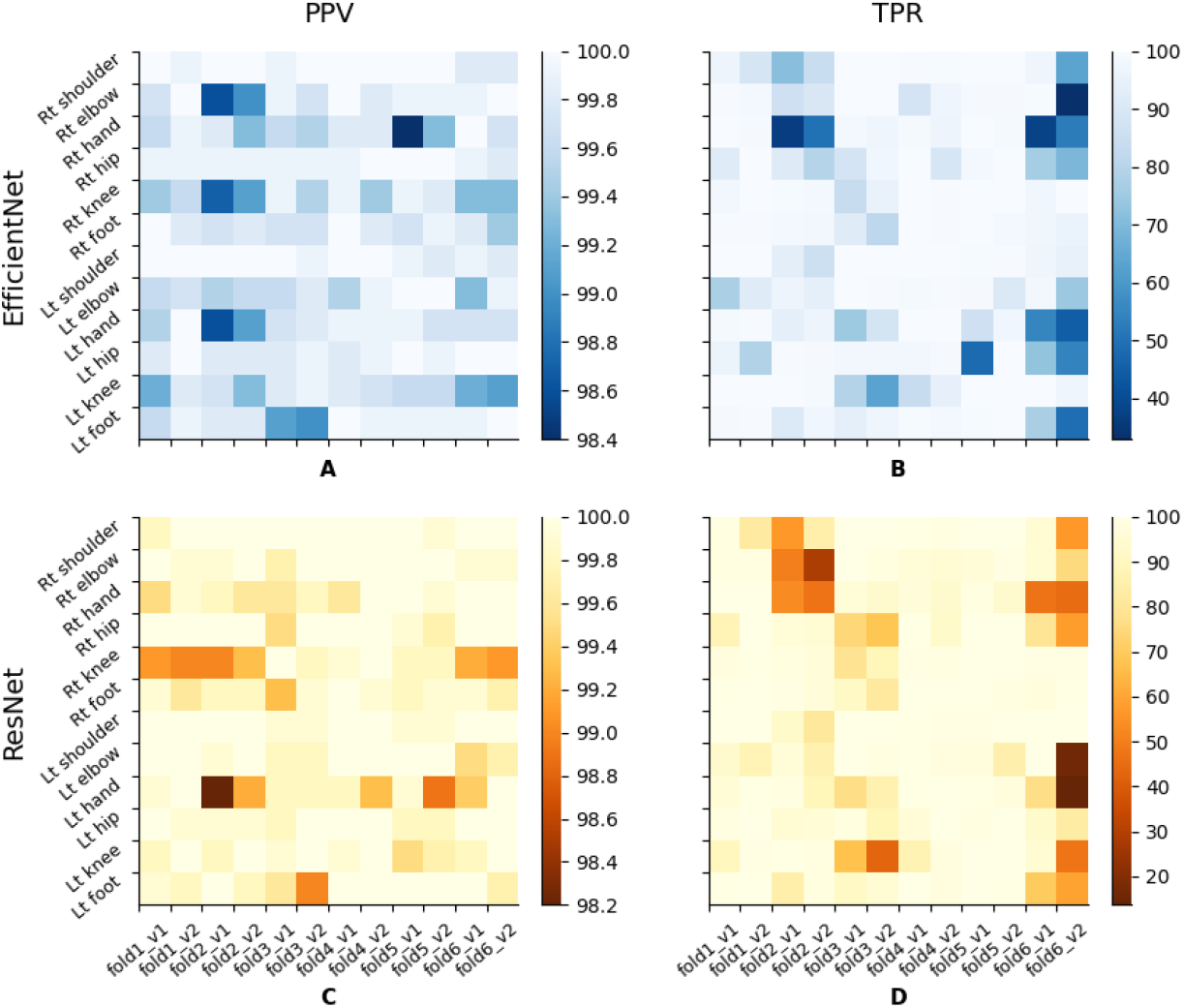
PPV and TPR measures across anatomical locations for each fold (unseen tested video) using efficientnet (A-B) and resnet (C-D).

**Figure 5.**
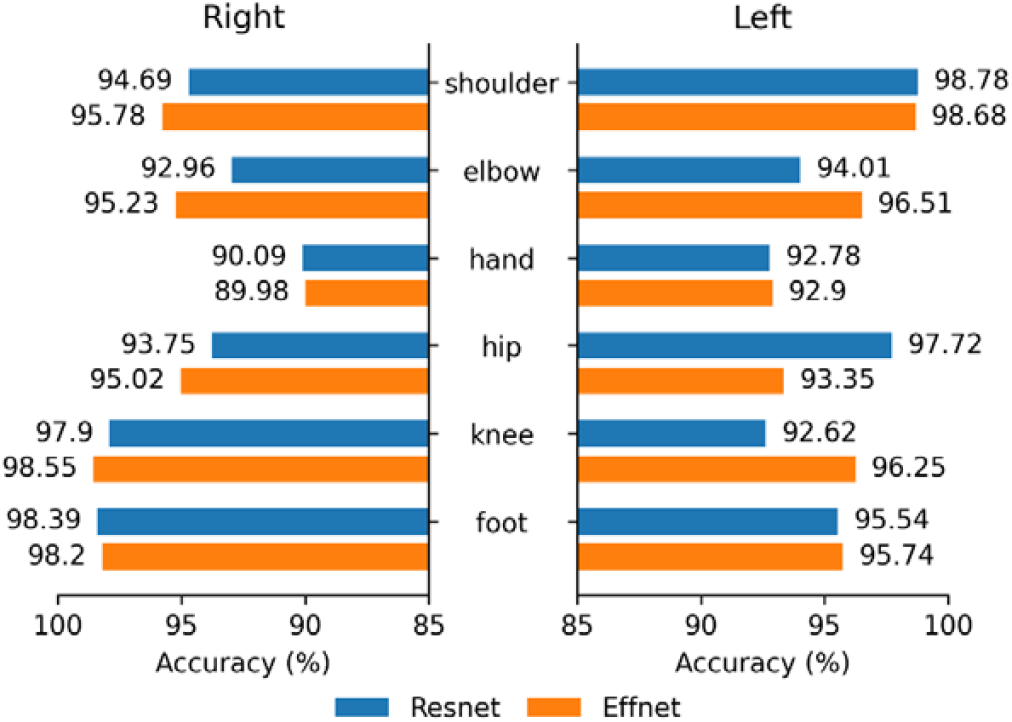
Location-based accuracy measures for the resnet-152 and efficientnet-b6.

Overall performance of the resnet and efficientnet for the subject-based and location-based approaches are shown in Fig. 5. The small performance variations both for the subject-based and location-based schemes in this figure, in particular, indicate generalization capability of the schemes. However, performance across subjects was dominated by one poorly performing outlier, fold6_v2 (see also Fig. 4) which will be discussed in the Discussion.

Examples of the algorithm’s predicted locations (in the test sets) versus manual annotations by the observer (HA) have been shown by crosses and dots in Fig. 6 (magnified representations from the left arms).

**Figure 6.**
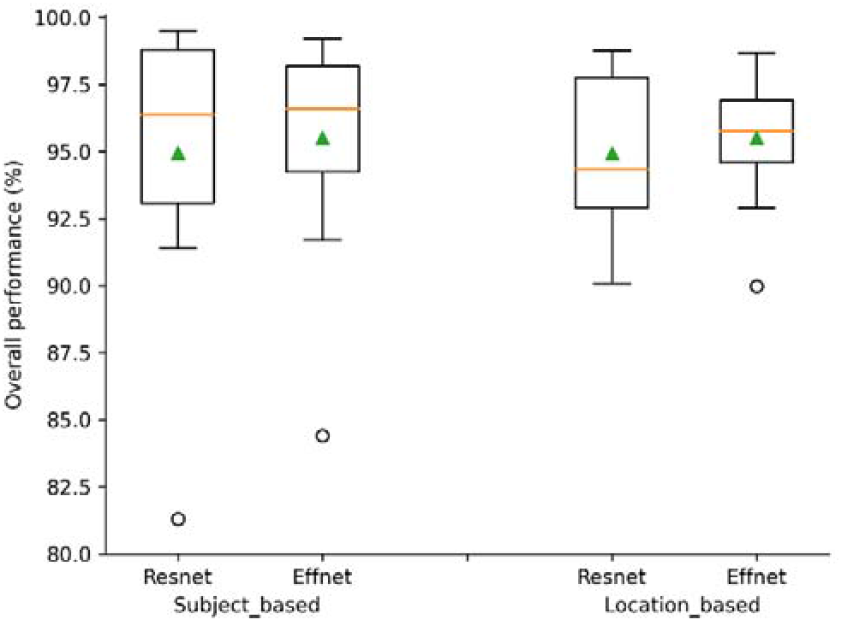
Overall accuracy of the subject-based and location-based schemes on the validation set. Green triangles represent mean values.

To further illustrate the performance of the deep-net classifiers, we visualized the precision (selectivity) and sensitivity measures for each of the classifiers across all folds for each video in each validation set. The boxplots in Fig 7 compare the precision and sensitivity measures separately for each individual video (v1 or v2) of the unseen infant using the resnet (5A and 5C) and the efficientnet (5B and 5D), respectively. A similar visualization approach was also carried out across all body-parts to illustrate what anatomical locations were associated with higher/lower precisions and sensitivities using each of the deep-net models, separately. The boxplots in Fig. S2 of the supplementary material section demonstrate these measures for each individual anatomical location across all folds using the resnet (8A and 8C) and the efficientnet (8B and 8D), respectively. Similarly, Fig. S3 demonstrates fold-based measurements across all anatomical locations.

**Figure 7.**
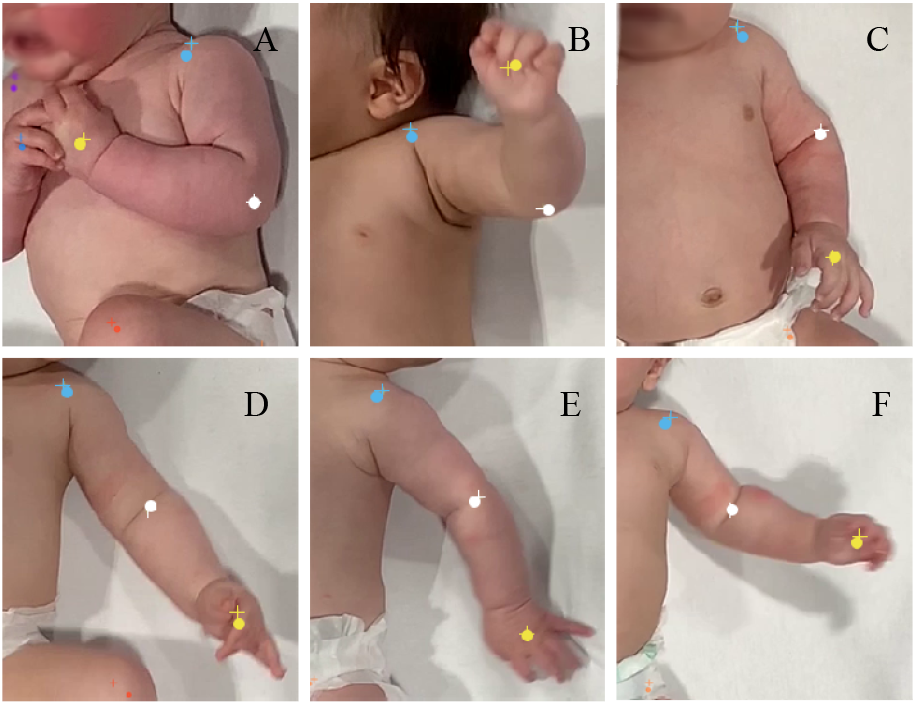
Examples of the automatic (+) vs manual (o) identifications in the left arms of 6 infants.

The scattered precision-recall (precision vs sensitivity) plots in Fig. S4 demonstrate how well the proposed deep-net models can track markers in each novel video of the unseen infant using data across 12 body-parts (subject-based scheme: resnet: A, efficientnet: C) and in each anatomical location across all tested videos (location-based scheme: resnet: B, efficientnet: D). An ideal precision-recall datapoint would be situated at the upper right corner which indicates higher precisions and sensitivities that are associated with fewer number of FPs and FNs respectively.

Comprehensive numerical details of the resnet and efficientnet models’ performance across all anatomical locations at each fold are represented in Tables S3-8 and S9-14 in the supporting information section, respectively. Figure 8 demonstrates sample trajectories extracted from the validation sets (unseen tested videos) using 200 frames per infant.

**Figure 8.**
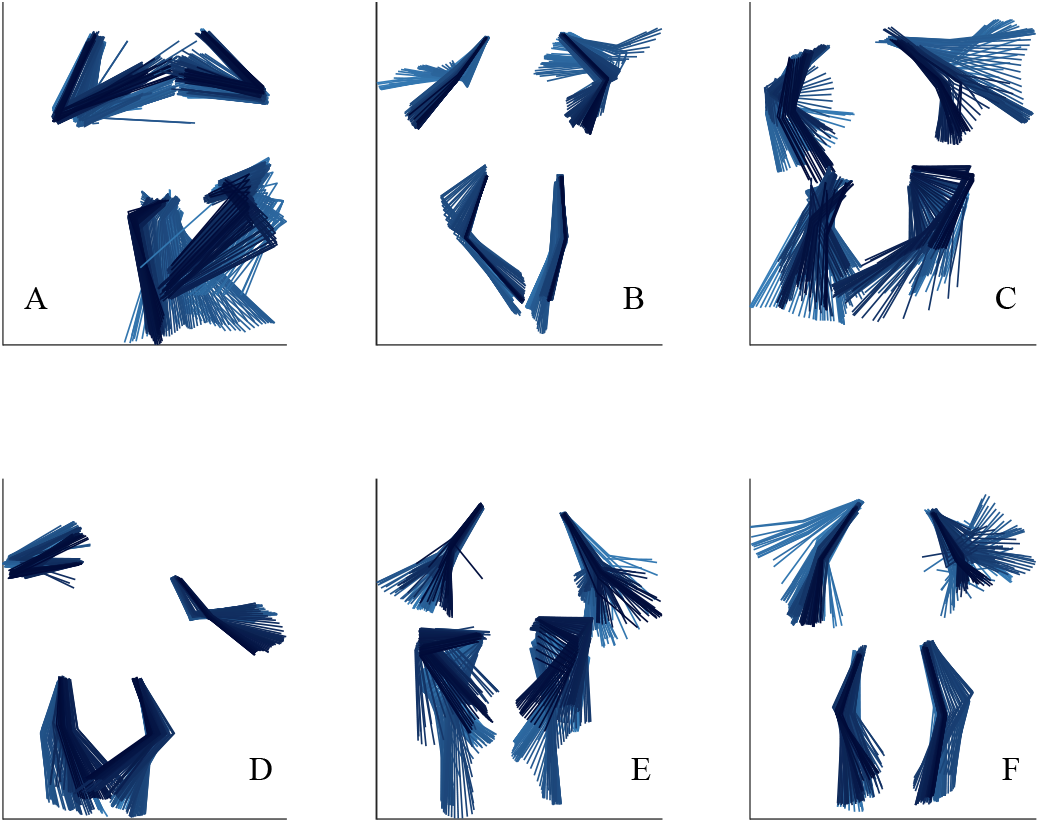
Example trajectories of 6.67 s (200 frames) length, extracted from the validation sets (unseen tested videos). Movements proceed from light to dark blue.

## 5. Discussion

This work proposes a novel markerless pose-estimation scheme, based on deep neural networks, for accurate motion tracking of infants’ movements using 2D video recordings from standard handheld devices (e.g., iPad). Subject-based performance assessment demonstrates generalization and performance consistency across out-of-domain data while identifying 12 different anatomical locations with overall cross-validated performances across 12 out-of-domain recordings from six infants of 95.52% (SD 2.43) and 94.94% (SD 2.63), respectively. The work further introduced a novel unsupervised performance evaluation strategy by combining Kalman filter likelihoods and deep-net probabilities to automatically measure performance metrics on larger out-of-domain datasets where manual labeling is challenging. Despite dealing with a relatively large validation set that includes ∼67k images, the performance range from this work resonates well with the performance range in recent markerless motion tracking studies performed on substantially smaller validations sets [39].

### Types of errors

Results indicate that despite postural variations in landmarks caused by rotational and/or fast movements in some body-parts, the proposed models have been able to extract and learn features related to each anatomical location in the 2D recordings of the training sets and later identify these locations correctly in the unseen data from novel infants (validation sets; see Tables S1 and S2). Further, validation of the models on data from novel infants was associated with consistently higher precision measures (corresponding to fewer FPs) compared to sensitivity measures (corresponding to more variable FN performance) across all folds. This result is important, mainly because it shows good generalization capability for the model obtained from a relatively small number of participants in the database, presumably aided by the relatively large number of frames used per participant [39].

### Effect of location

The location-based performance for the deep-net models ranged between 84.35-99.24% and 81.27 to 99.51% for the efficientnet and resnet, respectively across all anatomical locations (Tables S1 and S2). The lowest accuracy measures across both models were consistently associated with hands, which naturally exist in a much larger range of postures and speeds, and hips, which lack detailed, precisely located landmarks for the algorithm to identify (Figure 5). Patterns that were present near the hips were prints on the nappies of the infants, which varied in location and style between individuals and so were not useful for learning. Conversely, the left shoulder, right knee, and right foot are consistently associated with best tracking results.

The labeled hand poses in the training set from the five infants in each fold may not have fully covered the range of hand poses in the videos of the sixth infant in the validation sets. In that case, the algorithms would fail to identify a true observation due to lack of confidence (as they may not have seen a similar case before) and therefore, will label the observation as a false negative. This scenario was seen in the results for fold6_v2, where the baby had his hands locked into each other almost throughout the entire recording, resulting in poorer TPR performance of 45.95% and 45.10% for the resnet and efficientnet, respectively (see also Table S1 and S2). This hand posture had not been observed in the training set from the other five infants associated with this fold, who mostly kept their hands separated.

### Effect of velocity

We further investigated possible effects of velocity in hands and feet on the overall tracking performance. The tracking performance for feet markers was high despite the feet’s often varying dynamics (excess of movement and range) and speed (Table S1 and S2). Normalized histograms of the velocity of hands and feet across all 12 videos from all participants are shown in Figure S5. The close similarity between hand and feet velocity distributions suggests that the lower tracking performance of the hands is rather associated with hands’ natural morphological variations than the velocity itself: feet have a limited range of pose variations compared to hands, such that foot poses sampled in the training sets covered the range of potential poses, whereas this was not true for hands.

Fast movement exacerbates motion blur, where objects in the image move during the acquisition duration of individual frames. One mitigation of motion blur is to use video with a faster frame rate. Standard video frame rate is around 30 Hz, but newer consumer devices can record at 60 Hz or higher. The benefit for tracking accuracy of acquiring data at these higher rates is a useful avenue for future research.

### Effect of camera angle

A novel camera angle critically affected accuracy in our sample. Video fold6_v1 was filmed from quite a tilted angle on the left side compared to the other babies in the training set. The accuracies for this video were 84.35% and 81.27% across all folds for the efficientnet and resnet, respectively. In contrast, the overall performances improve to 91.73% and 93.62%, respectively, for the other video of the same infant movements from an overhead position (fold6_v2). In this overhead video, the true positive rate for the right hand still remains low due to the hands being locked into each other. For other infants, the proposed scheme was able to perform almost equally on the two videos despite smaller differences in viewing angle (e.g., fold 1 video 1: 98.04% vs video 2: 98.08% for the efficientnet, and fold 1 video 1: 98.49% vs video 2: 98.22% for the resnet).

### Other factors

Other factors that might have impaired body-part labelling include high-contrast shadows, which occurred in our images, infant’s skin color, and change of patterns on the nappies. Pre-processing of the videos to center the babies in their recordings, consistent use of a white background (mattress sheet), consistent lighting, and use of a fixed camera (as opposed to holding by hand) may have assisted with minimizing other sources of variation. These variables may be less well controlled when videos are recorded at home or in clinical settings.

Findings from this work indicate the feasibility of the proposed automated markerless movement tracking for reliable identification of infants’ movements in their 2D video recordings, for the purpose of the GMA. Our results overall support the premise that a rich training dataset covering the range of features expected in the novel data will enable a high level of generalization to new individuals. These results indicate that our approach is potentially suitable for use in a clinical platform for automating the GMA, which could be used alongside the current observational protocols to diagnose at-risk infants.

## Acknowledgements

- The research was supported by Friedlander Foundation grant (Grant: 3720759).
- The algorithm development, data analysis and manuscript writing/preparation were undertaken by H.A. Data was recorded by S.M. and L.L. Manuscript was reviewed and revised by A.M., S.W and T.B. Funding acquisition: T.B. and A.M. The final submitted article has been revised and approved by all authors.
- We would also like to acknowledge the use of New Zealand eScience Infrastructure (NeSI) high performance computing facilities to the results of this research. URL: https://www.nesi.org.nz.

## Supporting Information

**Table S1.** Resnet-152 overall performance (^*^marker-based assessment (or location based), ^**^ subject-based assessment). O. Perf.: overall performance=avgerage of PPV and TPR.

|  | *fold1_v1* |  | *fold1_v2* |  | *fold2_v1* |  | *fold2_v2* |  | *fold3_v1* |  | *fold3_v2* |  | *fold4_v1* |  | *fold4_v2* |  | *fold5_v1* |  | *fold5_v2* |  | *fold6_v1* |  | *fold6_v2* |  | *O. Perf.\** |
| --- | --- | --- | --- | --- | --- | --- | --- | --- | --- | --- | --- | --- | --- | --- | --- | --- | --- | --- | --- | --- | --- | --- | --- | --- | --- |
| *Total #frames* | 6558 |  | 6557 |  | 6818 |  | 6601 |  | 7456 |  | 7154 |  | 6352 |  | 6155 |  | 5953 |  | 5675 |  | 1934 |  | 1852 |  |  |
|  | *PPV* | *TPR* | *PPV* | *TPR* | *PPV* | *TPR* | *PPV* | *TPR* | *PPV* | *TPR* | *PPV* | *TPR* | *PPV* | *TPR* | *PPV* | *TPR* | *PPV* | *TPR* | *PPV* | *TPR* | *PPV* | *TPR* | *PPV* | *TPR* |  |
| *Rt shoulder* | 99.8 | 98.8 | 100 | 82.2 | 100 | 56.8 | 100 | 84.6 | 100 | 100 | 100 | 100 | 100 | 99.6 | 100 | 98.8 | 100 | 100 | 99.9 | 99.9 | 100 | 95.3 | 100 | 56.8 | 94.69 |
| *Rt elbow* | 100 | 98.8 | 99.9 | 98.9 | 99.9 | 50.0 | 100 | 28.7 | 99.7 | 100 | 100 | 98.3 | 100 | 96.6 | 100 | 94.8 | 100 | 96.1 | 100 | 99.2 | 99.9 | 95.4 | 99.9 | 75.2 | 92.96 |
| *Rt hand* | 99.5 | 99.9 | 99.9 | 99.8 | 99.8 | 53.2 | 99.6 | 46.8 | 99.6 | 96.4 | 99.8 | 93.6 | 99.6 | 96.8 | 100 | 93.9 | 100 | 98.7 | 99.9 | 93.4 | 100 | 47.0 | 100 | 44.9 | 90.09 |
| *Rt hip* | 100 | 87.8 | 100 | 100 | 100 | 96.6 | 100 | 95.4 | 99.5 | 74.1 | 100 | 68.0 | 100 | 99.5 | 100 | 92.9 | 99.9 | 99.8 | 99.7 | 100 | 100 | 79.3 | 100 | 57.6 | 93.75 |
| *Rt knee* | 99.1 | 97.7 | 99.0 | 99.5 | 99.0 | 98.1 | 99.3 | 96.7 | 100 | 78.6 | 99.8 | 88.6 | 99.9 | 99.3 | 100 | 98.8 | 99.8 | 100 | 99.8 | 100 | 99.2 | 99.2 | 99.1 | 99.3 | 97.90 |
| *Rt foot* | 99.9 | 99.4 | 99.6 | 99.8 | 99.8 | 99.1 | 99.8 | 96.8 | 99.3 | 91.4 | 99.9 | 81.7 | 100 | 100 | 99.9 | 100 | 99.8 | 100 | 99.9 | 98.6 | 99.9 | 97.6 | 99.7 | 99.4 | 98.39 |
| *Lt shoulder* | 100 | 99.9 | 100 | 99.9 | 100 | 92.7 | 100 | 81.0 | 99.9 | 100 | 99.9 | 100 | 100 | 99.6 | 100 | 99.0 | 99.9 | 99.3 | 99.9 | 99.7 | 100 | 100 | 100 | 100 | 98.78 |
| *Lt elbow* | 100 | 93.6 | 100 | 87.5 | 99.9 | 96.6 | 100 | 85.9 | 99.8 | 99.8 | 99.8 | 98.7 | 100 | 99.6 | 100 | 97.8 | 100 | 97.4 | 100 | 84.9 | 99.5 | 99.8 | 99.7 | 16.1 | 94.01 |
| *Lt hand* | 99.9 | 95.9 | 100 | 99.4 | 98.2 | 98.9 | 99.2 | 88.4 | 99.8 | 76.4 | 99.8 | 86.2 | 99.8 | 99.6 | 99.3 | 99.9 | 99.9 | 99.6 | 98.9 | 97.6 | 99.4 | 76.9 | 100 | 13.6 | 92.78 |
| *Lt hip* | 100 | 98.4 | 99.9 | 99.9 | 99.9 | 95.1 | 99.9 | 95.7 | 99.8 | 98.0 | 100 | 88.2 | 100 | 96.1 | 100 | 99.8 | 99.8 | 98.8 | 99.8 | 99.9 | 100 | 93.0 | 100 | 83.1 | 97.72 |
| *Lt knee* | 99.8 | 89.7 | 100 | 99.0 | 99.8 | 99.4 | 100 | 98.5 | 99.9 | 67.2 | 100 | 43.1 | 99.9 | 87.1 | 100 | 97.8 | 99.5 | 100 | 99.7 | 100 | 99.8 | 95.4 | 100 | 47.2 | 92.62 |
| *Lt foot* | 99.9 | 99.6 | 99.8 | 100 | 100 | 84.6 | 99.8 | 98.1 | 99.6 | 91.3 | 99.0 | 95.1 | 100 | 99.8 | 100 | 97.7 | 100 | 99.8 | 100 | 99.6 | 100 | 70.3 | 99.7 | 59.3 | 95.54 |
| *O. Perf. \*\** | 98.2 |  | 98.5 |  | 92.4 |  | 91.4 |  | 94.6 |  | 93.3 |  | 98.9 |  | 98.8 |  | 99.5 |  | 98.8 |  | 93.6 |  | 81.3 |  |  |

**Table S2.** Effnet-b6 overall performance (^*^marker-based assessment (or location based), ^**^ subject-based assessment). O. Perf.: overall performance=avgerage of PPV and TPR.

|  | fold1_v1 |  | fold1_v2 |  | fold2_v1 |  | fold2_v2 |  | fold3_v1 |  | fold3_v2 |  | fold4_v1 |  | fold4_v2 |  | fold5_v1 |  | fold5_v2 |  | fold6_v1 |  | fold6_v2 |  | O.Perf.* |
| --- | --- | --- | --- | --- | --- | --- | --- | --- | --- | --- | --- | --- | --- | --- | --- | --- | --- | --- | --- | --- | --- | --- | --- | --- | --- |
| Total #frames | 6558 |  | 6557 |  | 6818 |  | 6601 |  | 7456 |  | 7154 |  | 6352 |  | 6155 |  | 5953 |  | 5675 |  | 1934 |  | 1852 |  |  |
|  | PP<br>V | TP<br>R | PP<br>V | TP<br>R | PP<br>V | TP<br>R | PP<br>V | TP<br>R | PP<br>V | TP<br>R | PP<br>V | TP<br>R | PP<br>V | TP<br>R | PP<br>V | TP<br>R | PP<br>V | TP<br>R | PP<br>V | TP<br>R | PP<br>V | TP<br>R | PP<br>V | TP<br>R |  |
| Rt shoulder | 100 | 96.5 | 99.9 | 88.5 | 100 | 71.3 | 100 | 84.2 | 99.9 | 99.9 | 100 | 100 | 100 | 99.6 | 100 | 99.3 | 100 | 100 | 100 | 99.9 | 99.8 | 96.9 | 99.8 | 63.5 | 95.78 |
| Rt elbow | 99.7 | 99.5 | 100 | 98.4 | 98.6 | 86.0 | 99.0 | 89.7 | 99.9 | 100 | 99.7 | 100 | 100 | 88.1 | 99.8 | 96.8 | 99.9 | 99.2 | 99.9 | 99.2 | 99.9 | 99.4 | 100 | 32.9 | 95.23 |
| Rt hand | 99.6 | 100 | 99.9 | 99.1 | 99.8 | 36.8 | 99.3 | 49.9 | 99.6 | 98.3 | 99.5 | 96.9 | 99.8 | 98.8 | 99.8 | 96.4 | 98.4 | 99.9 | 99.3 | 98.7 | 100 | 37.7 | 99.7 | 52.5 | 89.98 |
| Rt hip | 99.9 | 92.1 | 99.9 | 100 | 99.9 | 91.6 | 99.9 | 79.7 | 99.9 | 88.1 | 99.9 | 97.4 | 99.9 | 99.8 | 100 | 89.0 | 100 | 98.5 | 100 | 99.9 | 99.9 | 76.2 | 99.8 | 69.3 | 95.02 |
| Rt knee | 99.4 | 97.7 | 99.6 | 99.8 | 98.7 | 99.4 | 99.1 | 98.6 | 99.9 | 83.6 | 99.5 | 95.2 | 99.9 | 99.8 | 99.4 | 99.6 | 99.9 | 100 | 99.8 | 100 | 99.3 | 97.5 | 99.3 | 99.9 | 98.55 |
| Rt foot | 100 | 99.5 | 99.8 | 99.7 | 99.7 | 98.5 | 99.8 | 97.7 | 99.7 | 91.7 | 99.7 | 81.3 | 100 | 100 | 99.8 | 99.6 | 99.7 | 100 | 99.9 | 98.6 | 99.8 | 97.3 | 99.4 | 95.6 | 98.20 |
| Lt shoulder | 100 | 100 | 100 | 100 | 100 | 93.1 | 100 | 85.8 | 100 | 100 | 99.9 | 100 | 100 | 99.8 | 100 | 100 | 99.9 | 99.3 | 99.8 | 99.7 | 99.9 | 96.6 | 99.8 | 94.9 | 98.68 |
| Lt elbow | 99.6 | 77.6 | 99.7 | 91.9 | 99.5 | 96.5 | 99.6 | 93.8 | 99.6 | 100 | 99.8 | 99.5 | 99.5 | 98.8 | 99.9 | 99.7 | 100 | 99.0 | 100 | 90.5 | 99.3 | 98.1 | 99.9 | 74.4 | 96.51 |
| Lt hand | 99.5 | 98.8 | 100 | 99.6 | 98.6 | 93.8 | 99.1 | 96.4 | 99.7 | 74.8 | 99.8 | 87.7 | 99.9 | 99.8 | 99.9 | 99.9 | 99.9 | 85.9 | 99.7 | 97.6 | 99.7 | 54.9 | 99.7 | 44.9 | 92.90 |
| Lt hip | 99.8 | 96.1 | 100 | 78.9 | 99.8 | 99.6 | 99.8 | 99.0 | 99.8 | 97.9 | 99.9 | 98.0 | 99.8 | 97.8 | 99.9 | 99.1 | 100 | 48.2 | 99.9 | 99.9 | 100 | 72.8 | 100 | 54.3 | 93.35 |
| Lt knee | 99.2 | 99.8 | 99.8 | 100 | 99.7 | 99.9 | 99.3 | 98.8 | 99.8 | 79.3 | 99.9 | 62.9 | 99.8 | 83.9 | 99.7 | 94.3 | 99.6 | 100 | 99.6 | 100 | 99.2 | 99.9 | 99.1 | 96.6 | 96.25 |
| Lt foot | 99.6 | 99.2 | 99.9 | 99.6 | 99.8 | 90.7 | 99.8 | 97.5 | 99.1 | 93.5 | 99.0 | 96.8 | 100 | 99.4 | 99.9 | 98.5 | 99.9 | 99.4 | 99.9 | 99.8 | 99.9 | 77.4 | 100 | 49.0 | 95.74 |
| O. Perf.** | 98.0 |  | 98.1 |  | 93.8 |  | 94.4 |  | 96.0 |  | 96.3 |  | 98.5 |  | 98.8 |  | 96.9 |  | 99.2 |  | 91.7 |  | 84.4 |  |  |

**Figure S1.**
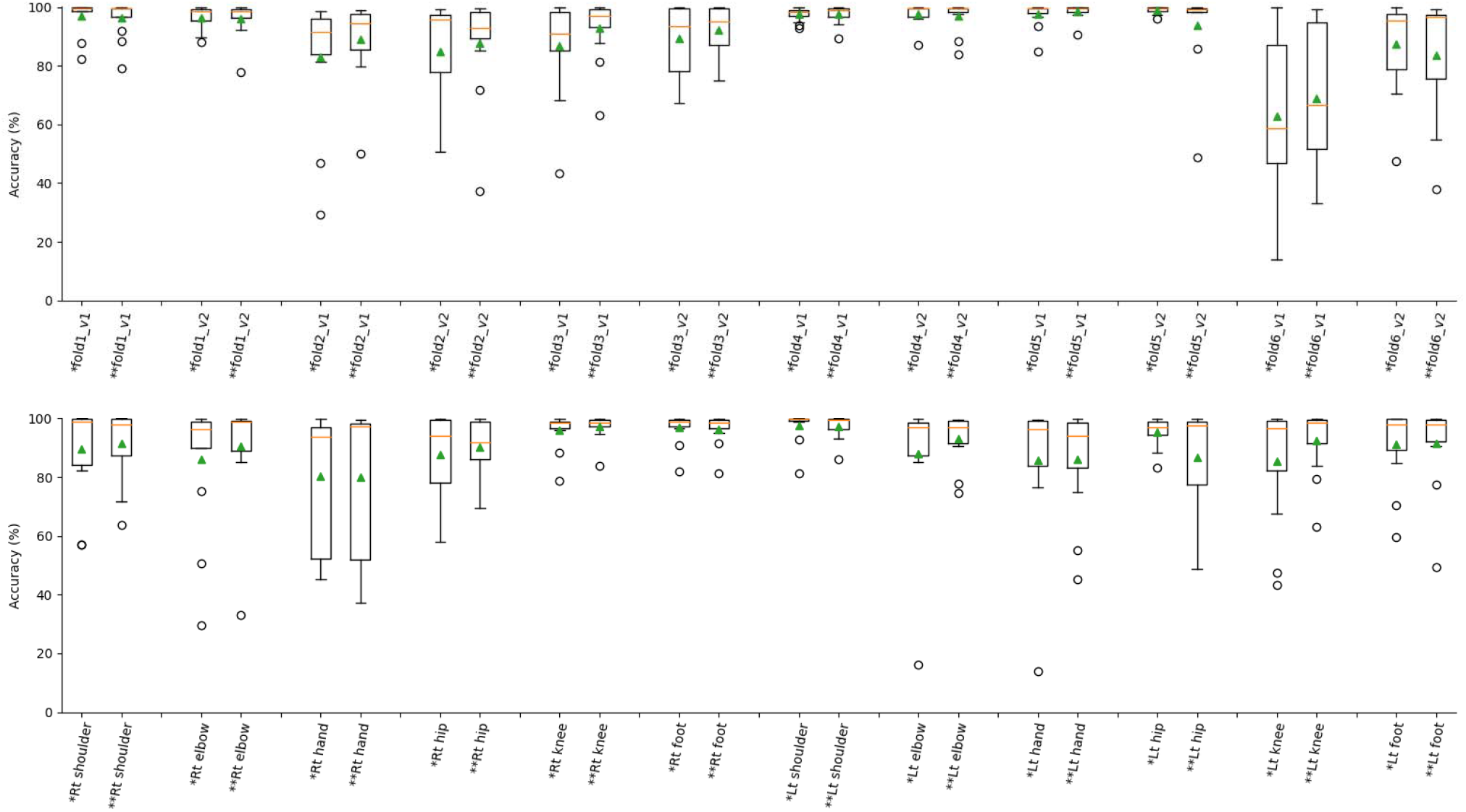
Detailed comparison schematic for the performance of the resnet(*) vs efficientnet(**) models in the subject-based (A) vs. location-based (B) schemes. Green triangles represent mean values.

### Resnet-152 results

**Table S3.** Fold_1, trained on data from infants # 1, 2, 3, 4, 5;

|  | *Test results on video #2 of baby 6* |  |  |  |  |  |  | *Test results on video #1 of baby 6* |  |  |  |  |  |  |
| --- | --- | --- | --- | --- | --- | --- | --- | --- | --- | --- | --- | --- | --- | --- |
|  | *TP* | *FP* | *FN* | *TN* | *Selectivity* | *Sensitivity* | *Accuracy* | *TP* | *FP* | *FN* | *TN* | *Selectivity* | *Sensitivity* | *Accuracy* |
| *Rt shoulder* | 5371 | 0 | 1162 | 25 | 100.00 | 82.21 | 82.28 | 6467 | 10 | 79 | 1 | 99.85 | 98.79 | 98.64 |
| *Rt elbow* | 6464 | 6 | 75 | 12 | 99.91 | 98.85 | 98.76 | 6452 | 0 | 81 | 24 | 100.00 | 98.76 | 98.76 |
| *Rt hand* | 6534 | 5 | 15 | 4 | 99.92 | 99.77 | 99.70 | 6521 | 32 | 4 | 0 | 99.51 | 99.94 | 99.45 |
| *Rt hip* | 6555 | 3 | 0 | 0 | 99.95 | 100.00 | 99.95 | 5712 | 1 | 793 | 51 | 99.98 | 87.81 | 87.89 |
| *Rt knee* | 6452 | 68 | 35 | 3 | 98.96 | 99.46 | 98.43 | 6317 | 60 | 151 | 29 | 99.06 | 97.67 | 96.78 |
| *Rt foot* | 6521 | 23 | 14 | 0 | 99.65 | 99.79 | 99.44 | 6512 | 4 | 40 | 1 | 99.94 | 99.39 | 99.33 |
| *Lt shoulder* | 6550 | 2 | 6 | 0 | 99.97 | 99.91 | 99.88 | 6548 | 3 | 4 | 2 | 99.95 | 99.94 | 99.89 |
| *Lt elbow* | 5677 | 1 | 813 | 89 | 99.98 | 87.47 | 87.63 | 6081 | 3 | 419 | 54 | 99.95 | 93.55 | 93.56 |
| *Lt hand* | 6502 | 2 | 40 | 14 | 99.97 | 99.39 | 99.36 | 6211 | 6 | 268 | 72 | 99.90 | 95.86 | 95.82 |
| *Lt hip* | 6540 | 7 | 7 | 4 | 99.89 | 99.89 | 99.79 | 6433 | 0 | 102 | 22 | 100.00 | 98.44 | 98.44 |
| *Lt knee* | 6486 | 2 | 68 | 2 | 99.97 | 98.96 | 98.93 | 5831 | 10 | 667 | 49 | 99.83 | 89.74 | 89.68 |
| *Lt foot* | 6540 | 12 | 3 | 3 | 99.82 | 99.95 | 99.77 | 6514 | 8 | 29 | 6 | 99.88 | 99.56 | 99.44 |
| *Overall* |  |  |  |  | 99.83 | 97.14 | 96.99 |  |  |  |  | 99.82 | 96.62 | 96.47 |

**Table S4.** Fold_2, trained on data from infants # 1, 2, 3, 4, 6;

|  | **Test results on video #2 of baby 5** |  |  |  |  |  |  | **Test results on video #1 of baby 5** |  |  |  |  |  |  |
| --- | --- | --- | --- | --- | --- | --- | --- | --- | --- | --- | --- | --- | --- | --- |
|  | *TP* | *FP* | *FN* | *TN* | *Selectivity* | *Sensitivity* | *Accuracy* | *TP* | *FP* | *FN* | *TN* | *Selectivity* | *Sensitivity* | *Accuracy* |
| *Rt shoulder* | 5728 | 0 | 1044 | 46 | 100.00 | 84.58 | 84.69 | 3711 | 0 | 2828 | 62 | 100.00 | 56.75 | 57.16 |
| *Rt elbow* | 1940 | 0 | 4813 | 65 | 100.00 | 28.73 | 29.41 | 3234 | 2 | 3234 | 80 | 99.94 | 50.00 | 50.60 |
| *Rt hand* | 3169 | 14 | 3601 | 34 | 99.56 | 46.81 | 46.98 | 3473 | 6 | 3057 | 64 | 99.83 | 53.19 | 53.59 |
| *Rt hip* | 6495 | 0 | 311 | 12 | 100.00 | 95.43 | 95.44 | 6352 | 0 | 222 | 27 | 100.00 | 96.62 | 96.64 |
| *Rt knee* | 6536 | 46 | 226 | 10 | 99.30 | 96.66 | 96.01 | 6404 | 62 | 122 | 11 | 99.04 | 98.13 | 97.21 |
| *Rt foot* | 6549 | 14 | 215 | 40 | 99.79 | 96.82 | 96.64 | 6510 | 15 | 62 | 14 | 99.77 | 99.06 | 98.83 |
| *Lt shoulder* | 5460 | 0 | 1281 | 77 | 100.00 | 81.00 | 81.21 | 6084 | 0 | 476 | 41 | 100.00 | 92.74 | 92.79 |
| *Lt elbow* | 5786 | 0 | 948 | 84 | 100.00 | 85.92 | 86.10 | 6332 | 6 | 223 | 40 | 99.91 | 96.60 | 96.53 |
| *Lt hand* | 5962 | 50 | 779 | 27 | 99.17 | 88.44 | 87.84 | 6412 | 115 | 70 | 4 | 98.24 | 98.92 | 97.20 |
| *Lt hip* | 6497 | 4 | 291 | 26 | 99.94 | 95.71 | 95.67 | 6261 | 7 | 326 | 7 | 99.89 | 95.05 | 94.96 |
| *Lt knee* | 6701 | 1 | 104 | 12 | 99.99 | 98.47 | 98.46 | 6543 | 13 | 37 | 8 | 99.80 | 99.44 | 99.24 |
| *Lt foot* | 6672 | 15 | 126 | 5 | 99.78 | 98.15 | 97.93 | 5540 | 2 | 1009 | 49 | 99.96 | 84.59 | 84.68 |
| **Overall** |  |  |  |  | 99.79 | 83.06 | 83.03 |  |  |  |  | 99.70 | 85.09 | 84.95 |

**Table S5.**
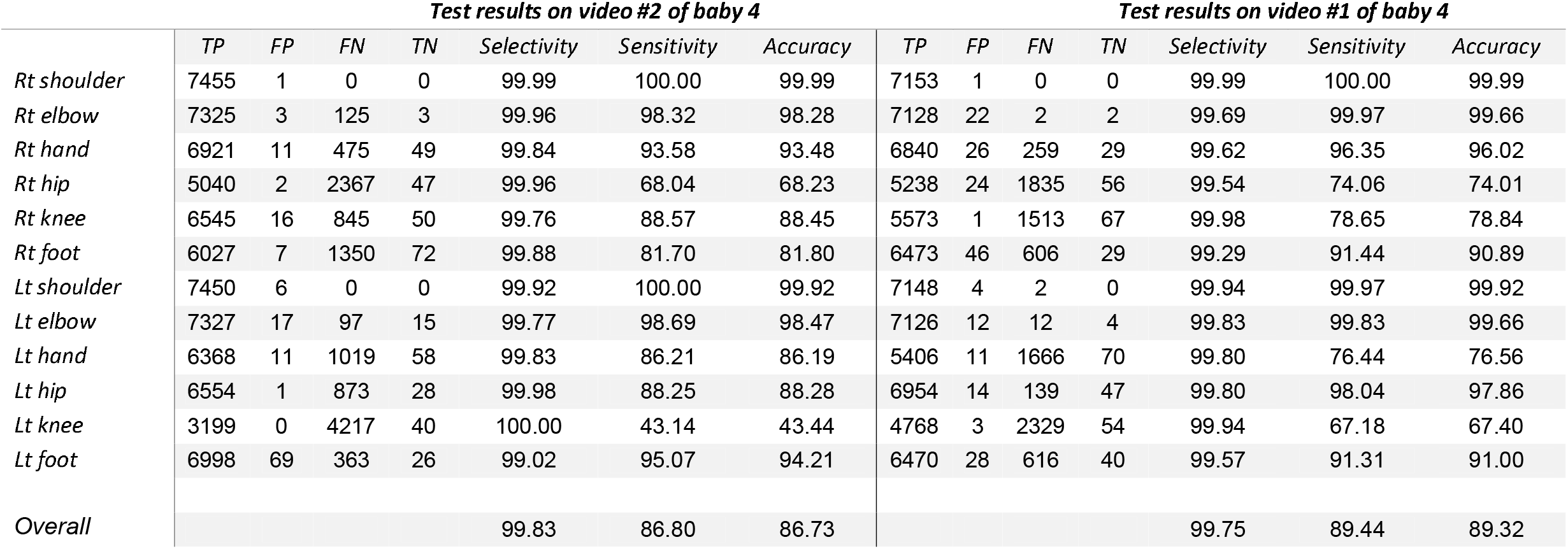
Fold_3, trained on data from infants # 1, 2, 3, 5, 6;

|  | *Test results on video #2 of baby 4* |  |  |  |  |  |  | *Test results on video #1 of baby 4* |  |  |  |  |  |  |
| --- | --- | --- | --- | --- | --- | --- | --- | --- | --- | --- | --- | --- | --- | --- |
|  | *TP* | *FP* | *FN* | *TN* | *Selectivity* | *Sensitivity* | *Accuracy* | *TP* | *FP* | *FN* | *TN* | *Selectivity* | *Sensitivity* | *Accuracy* |
| *Rt shoulder* | 7455 | 1 | 0 | 0 | 99.99 | 100.00 | 99.99 | 7153 | 1 | 0 | 0 | 99.99 | 100.00 | 99.99 |
| *Rt elbow* | 7325 | 3 | 125 | 3 | 99.96 | 98.32 | 98.28 | 7128 | 22 | 2 | 2 | 99.69 | 99.97 | 99.66 |
| *Rt hand* | 6921 | 11 | 475 | 49 | 99.84 | 93.58 | 93.48 | 6840 | 26 | 259 | 29 | 99.62 | 96.35 | 96.02 |
| *Rt hip* | 5040 | 2 | 2367 | 47 | 99.96 | 68.04 | 68.23 | 5238 | 24 | 1835 | 56 | 99.54 | 74.06 | 74.01 |
| *Rt knee* | 6545 | 16 | 845 | 50 | 99.76 | 88.57 | 88.45 | 5573 | 1 | 1513 | 67 | 99.98 | 78.65 | 78.84 |
| *Rt foot* | 6027 | 7 | 1350 | 72 | 99.88 | 81.70 | 81.80 | 6473 | 46 | 606 | 29 | 99.29 | 91.44 | 90.89 |
| *Lt shoulder* | 7450 | 6 | 0 | 0 | 99.92 | 100.00 | 99.92 | 7148 | 4 | 2 | 0 | 99.94 | 99.97 | 99.92 |
| *Lt elbow* | 7327 | 17 | 97 | 15 | 99.77 | 98.69 | 98.47 | 7126 | 12 | 12 | 4 | 99.83 | 99.83 | 99.66 |
| *Lt hand* | 6368 | 11 | 1019 | 58 | 99.83 | 86.21 | 86.19 | 5406 | 11 | 1666 | 70 | 99.80 | 76.44 | 76.56 |
| *Lt hip* | 6554 | 1 | 873 | 28 | 99.98 | 88.25 | 88.28 | 6954 | 14 | 139 | 47 | 99.80 | 98.04 | 97.86 |
| *Lt knee* | 3199 | 0 | 4217 | 40 | 100.00 | 43.14 | 43.44 | 4768 | 3 | 2329 | 54 | 99.94 | 67.18 | 67.40 |
| *Lt foot* | 6998 | 69 | 363 | 26 | 99.02 | 95.07 | 94.21 | 6470 | 28 | 616 | 40 | 99.57 | 91.31 | 91.00 |
| *Overall* |  |  |  |  | 99.83 | 86.80 | 86.73 |  |  |  |  | 99.75 | 89.44 | 89.32 |

**Table S6.** Fold_4, trained on data from infants # 1, 2, 4, 4, 6;

|  | **Test results on video #2 of baby 3** |  |  |  |  |  |  | **Test results on video #1 of baby 3** |  |  |  |  |  |  |
| --- | --- | --- | --- | --- | --- | --- | --- | --- | --- | --- | --- | --- | --- | --- |
|  | *TP* | *FP* | *FN* | *TN* | *Selectivity* | *Sensitivity* | *Accuracy* | *TP* | *FP* | *FN* | *TN* | *Selectivity* | *Sensitivity* | *Accuracy* |
| *Rt shoulder* | 6271 | 1 | 75 | 5 | 99.98 | 98.82 | 98.80 | 6126 | 1 | 24 | 4 | 99.98 | 99.61 | 99.59 |
| *Rt elbow* | 5994 | 1 | 328 | 29 | 99.98 | 94.81 | 94.82 | 5923 | 0 | 211 | 21 | 100.00 | 96.56 | 96.57 |
| *Rt hand* | 5941 | 0 | 387 | 24 | 100.00 | 93.88 | 93.91 | 5915 | 21 | 197 | 22 | 99.65 | 96.78 | 96.46 |
| *Rt hip* | 5866 | 0 | 450 | 36 | 100.00 | 92.88 | 92.92 | 6123 | 1 | 31 | 0 | 99.98 | 99.50 | 99.48 |
| *Rt knee* | 6262 | 0 | 77 | 13 | 100.00 | 98.79 | 98.79 | 6104 | 6 | 43 | 2 | 99.90 | 99.30 | 99.20 |
| *Rt foot* | 6347 | 5 | 0 | 0 | 99.92 | 100.00 | 99.92 | 6151 | 3 | 0 | 1 | 99.95 | 100.00 | 99.95 |
| *Lt shoulder* | 6283 | 0 | 66 | 3 | 100.00 | 98.96 | 98.96 | 6120 | 2 | 27 | 6 | 99.97 | 99.56 | 99.53 |
| *Lt elbow* | 6197 | 3 | 137 | 14 | 99.95 | 97.84 | 97.80 | 6110 | 0 | 22 | 23 | 100.00 | 99.64 | 99.64 |
| *Lt hand* | 6305 | 43 | 4 | 0 | 99.32 | 99.94 | 99.26 | 6115 | 15 | 22 | 3 | 99.76 | 99.64 | 99.40 |
| *Lt hip* | 6338 | 0 | 10 | 4 | 100.00 | 99.84 | 99.84 | 5859 | 1 | 240 | 55 | 99.98 | 96.06 | 96.08 |
| *Lt knee* | 6184 | 0 | 138 | 30 | 100.00 | 97.82 | 97.83 | 5288 | 6 | 786 | 75 | 99.89 | 87.06 | 87.13 |
| *Lt foot* | 6193 | 0 | 145 | 14 | 100.00 | 97.71 | 97.72 | 6143 | 1 | 11 | 0 | 99.98 | 99.82 | 99.81 |
| **Overall** |  |  |  |  | 99.93 | 97.61 | 97.55 |  |  |  |  | 99.92 | 97.79 | 97.74 |

**Table S7.** Fold_5, trained on data from infants # 1, 3, 4, 5, 6;

|  | *Test results on video #2 of baby 2* |  |  |  |  |  |  | *Test results on video #1 of baby 2* |  |  |  |  |  |  |
| --- | --- | --- | --- | --- | --- | --- | --- | --- | --- | --- | --- | --- | --- | --- |
|  | *TP* | *FP* | *FN* | *TN* | *Selectivity* | *Sensitivity* | *Accuracy* | *TP* | *FP* | *FN* | *TN* | *Selectivity* | *Sensitivity* | *Accuracy* |
| *Rt shoulder* | 5940 | 3 | 7 | 3 | 99.95 | 99.88 | 99.83 | 5672 | 1 | 0 | 2 | 99.98 | 100.00 | 99.98 |
| *Rt elbow* | 5894 | 0 | 46 | 13 | 100.00 | 99.23 | 99.23 | 5423 | 0 | 223 | 29 | 100.00 | 96.05 | 96.07 |
| *Rt hand* | 5540 | 4 | 389 | 20 | 99.93 | 93.44 | 93.40 | 5594 | 0 | 71 | 10 | 100.00 | 98.75 | 98.75 |
| *Rt hip* | 5937 | 16 | 0 | 0 | 99.73 | 100.00 | 99.73 | 5658 | 5 | 11 | 1 | 99.91 | 99.81 | 99.72 |
| *Rt knee* | 5939 | 14 | 0 | 0 | 99.76 | 100.00 | 99.76 | 5665 | 10 | 0 | 0 | 99.82 | 100.00 | 99.82 |
| *Rt foot* | 5854 | 3 | 84 | 12 | 99.95 | 98.59 | 98.54 | 5663 | 10 | 2 | 0 | 99.82 | 99.96 | 99.79 |
| *Lt shoulder* | 5925 | 6 | 19 | 3 | 99.90 | 99.68 | 99.58 | 5627 | 4 | 41 | 3 | 99.93 | 99.28 | 99.21 |
| *Lt elbow* | 5012 | 0 | 894 | 47 | 100.00 | 84.86 | 84.98 | 5498 | 0 | 149 | 28 | 100.00 | 97.36 | 97.37 |
| *Lt hand* | 5721 | 61 | 138 | 33 | 98.95 | 97.64 | 96.66 | 5642 | 6 | 25 | 2 | 99.89 | 99.56 | 99.45 |
| *Lt hip* | 5932 | 13 | 6 | 2 | 99.78 | 99.90 | 99.68 | 5597 | 10 | 66 | 2 | 99.82 | 98.83 | 98.66 |
| *Lt knee* | 5933 | 20 | 0 | 0 | 99.66 | 100.00 | 99.66 | 5648 | 27 | 0 | 0 | 99.52 | 100.00 | 99.52 |
| *Lt foot* | 5927 | 0 | 23 | 3 | 100.00 | 99.61 | 99.61 | 5663 | 1 | 9 | 2 | 99.98 | 99.84 | 99.82 |
| *Overall* |  |  |  |  | 99.80 | 97.74 | 97.56 |  |  |  |  | 99.89 | 99.12 | 99.01 |

**Table S8.** Fold_6, trained on data from infants # 2, 3, 4, 5, 6;

|  | *Test results on video #2 of baby 1* |  |  |  |  |  |  | *Test results on video #1 of baby 1* |  |  |  |  |  |  |
| --- | --- | --- | --- | --- | --- | --- | --- | --- | --- | --- | --- | --- | --- | --- |
|  | *TP* | *FP* | *FN* | *TN* | *Selectivity* | *Sensitivity* | *Accuracy* | *TP* | *FP* | *FN* | *TN* | *Selectivity* | *Sensitivity* | *Accuracy* |
| *Rt shoulder* | 1089 | 0 | 829 | 16 | 100.00 | 56.78 | 57.14 | 1762 | 0 | 87 | 3 | 100.00 | 95.29 | 95.30 |
| *Rt elbow* | 1441 | 1 | 476 | 16 | 99.93 | 75.17 | 75.34 | 1754 | 2 | 85 | 11 | 99.89 | 95.38 | 95.30 |
| *Rt hand* | 861 | 0 | 1058 | 15 | 100.00 | 44.87 | 45.29 | 861 | 0 | 972 | 19 | 100.00 | 46.97 | 47.52 |
| *Rt hip* | 1107 | 0 | 815 | 12 | 100.00 | 57.60 | 57.86 | 1456 | 0 | 381 | 15 | 100.00 | 79.26 | 79.43 |
| *Rt knee* | 1902 | 18 | 13 | 1 | 99.06 | 99.32 | 98.40 | 1821 | 14 | 15 | 2 | 99.24 | 99.18 | 98.43 |
| *Rt foot* | 1913 | 6 | 12 | 3 | 99.69 | 99.38 | 99.07 | 1798 | 2 | 44 | 8 | 99.89 | 97.61 | 97.52 |
| *Lt shoulder* | 1934 | 0 | 0 | 0 | 100.00 | 100.00 | 100.00 | 1852 | 0 | 0 | 0 | 100.00 | 100.00 | 100.00 |
| *Lt elbow* | 310 | 1 | 1618 | 3 | 99.68 | 16.08 | 16.20 | 1839 | 10 | 3 | 0 | 99.46 | 99.84 | 99.30 |
| *Lt hand* | 262 | 0 | 1666 | 6 | 100.00 | 13.59 | 13.86 | 1407 | 9 | 423 | 13 | 99.36 | 76.89 | 76.67 |
| *Lt hip* | 1593 | 0 | 323 | 18 | 100.00 | 83.14 | 83.30 | 1720 | 0 | 129 | 3 | 100.00 | 93.02 | 93.03 |
| *Lt knee* | 909 | 0 | 1017 | 8 | 100.00 | 47.20 | 47.41 | 1749 | 3 | 84 | 16 | 99.83 | 95.42 | 95.30 |
| *Lt foot* | 1139 | 3 | 782 | 10 | 99.74 | 59.29 | 59.41 | 1293 | 0 | 545 | 14 | 100.00 | 70.35 | 70.57 |
| *Overall* |  |  |  |  | 99.84 | 62.70 | 62.77 |  |  |  |  | 99.81 | 87.43 | 87.37 |

### Efficientnet-b6 results

**Table S9.** Fold_1, trained on data from infants # 1, 2, 3, 4, 5;

|  | *Test results on video #2 of baby 6* |  |  |  |  |  |  | *Test results on video #1 of baby 6* |  |  |  |  |  |  |
| --- | --- | --- | --- | --- | --- | --- | --- | --- | --- | --- | --- | --- | --- | --- |
|  | *TP* | *FP* | *FN* | *TN* | *Selectivity* | *Sensitivity* | *Accuracy* | *TP* | *FP* | *FN* | *TN* | *Selectivity* | *Sensitivity* | *Accuracy* |
| *Rt shoulder* | 5756 | 8 | 751 | 43 | 99.86 | 88.46 | 88.43 | 6306 | 0 | 226 | 25 | 100.00 | 96.54 | 96.55 |
| *Rt elbow* | 6415 | 1 | 107 | 35 | 99.98 | 98.36 | 98.35 | 6464 | 17 | 34 | 42 | 99.74 | 99.48 | 99.22 |
| *Rt hand* | 6471 | 7 | 62 | 18 | 99.89 | 99.05 | 98.95 | 6528 | 27 | 2 | 0 | 99.59 | 99.97 | 99.56 |
| *Rt hip* | 6553 | 5 | 0 | 0 | 99.92 | 100.00 | 99.92 | 5959 | 5 | 510 | 83 | 99.92 | 92.12 | 92.15 |
| *Rt knee* | 6511 | 28 | 10 | 9 | 99.57 | 99.85 | 99.42 | 6312 | 39 | 147 | 59 | 99.39 | 97.72 | 97.16 |
| *Rt foot* | 6525 | 10 | 18 | 5 | 99.85 | 99.72 | 99.57 | 6506 | 3 | 34 | 14 | 99.95 | 99.48 | 99.44 |
| *Lt shoulder* | 6557 | 0 | 1 | 0 | 100.00 | 99.98 | 99.98 | 6551 | 2 | 2 | 2 | 99.97 | 99.97 | 99.94 |
| *Lt elbow* | 5929 | 18 | 525 | 86 | 99.70 | 91.87 | 91.72 | 4984 | 18 | 1440 | 115 | 99.64 | 77.58 | 77.76 |
| *Lt hand* | 6516 | 2 | 24 | 16 | 99.97 | 99.63 | 99.60 | 6430 | 31 | 77 | 19 | 99.52 | 98.82 | 98.35 |
| *Lt hip* | 5122 | 2 | 1372 | 62 | 99.96 | 78.87 | 79.05 | 6217 | 11 | 254 | 75 | 99.82 | 96.07 | 95.96 |
| *Lt knee* | 6544 | 14 | 0 | 0 | 99.79 | 100.00 | 99.79 | 6492 | 52 | 11 | 2 | 99.21 | 99.83 | 99.04 |
| *Lt foot* | 6524 | 7 | 23 | 4 | 99.89 | 99.65 | 99.54 | 6469 | 25 | 55 | 8 | 99.62 | 99.16 | 98.78 |
| *Overall* |  |  |  |  | 99.87 | 96.29 | 96.19 |  |  |  |  | 99.70 | 96.39 | 96.16 |

**Table S10.** Fold_2, trained on data from infants # 1, 2, 3, 4, 6;

|  | **Test results on video #2 of baby 5** |  |  |  |  |  |  | **Test results on video #1 of baby 5** |  |  |  |  |  |  |
| --- | --- | --- | --- | --- | --- | --- | --- | --- | --- | --- | --- | --- | --- | --- |
|  | *TP* | *FP* | *FN* | *TN* | *Selectivity* | *Sensitivity* | *Accuracy* | *TP* | *FP* | *FN* | *TN* | *Selectivity* | *Sensitivity* | *Accuracy* |
| *Rt shoulder* | 5689 | 2 | 1070 | 57 | 99.96 | 84.17 | 84.28 | 4643 | 2 | 1870 | 86 | 99.96 | 71.29 | 71.64 |
| *Rt elbow* | 6034 | 58 | 693 | 33 | 99.05 | 89.70 | 88.99 | 5575 | 79 | 909 | 38 | 98.60 | 85.98 | 85.03 |
| *Rt hand* | 3374 | 24 | 3389 | 31 | 99.29 | 49.89 | 49.94 | 2414 | 5 | 4141 | 41 | 99.79 | 36.83 | 37.19 |
| *Rt hip* | 5357 | 5 | 1367 | 89 | 99.91 | 79.67 | 79.88 | 5959 | 5 | 546 | 91 | 99.92 | 91.61 | 91.65 |
| *Rt knee* | 6643 | 57 | 97 | 21 | 99.15 | 98.56 | 97.74 | 6461 | 85 | 41 | 14 | 98.70 | 99.37 | 98.09 |
| *Rt foot* | 6634 | 16 | 153 | 15 | 99.76 | 97.75 | 97.52 | 6478 | 20 | 97 | 6 | 99.69 | 98.52 | 98.23 |
| *Lt shoulder* | 5776 | 1 | 953 | 88 | 99.98 | 85.84 | 86.01 | 6076 | 0 | 451 | 74 | 100.00 | 93.09 | 93.17 |
| *Lt elbow* | 6331 | 23 | 421 | 63 | 99.64 | 93.76 | 93.51 | 6294 | 33 | 227 | 47 | 99.48 | 96.52 | 96.06 |
| *Lt hand* | 6498 | 60 | 246 | 14 | 99.09 | 96.35 | 95.51 | 6110 | 89 | 401 | 1 | 98.56 | 93.84 | 92.58 |
| *Lt hip* | 6713 | 12 | 68 | 25 | 99.82 | 99.00 | 98.83 | 6551 | 10 | 26 | 14 | 99.85 | 99.60 | 99.45 |
| *Lt knee* | 6664 | 50 | 83 | 21 | 99.26 | 98.77 | 98.05 | 6570 | 20 | 4 | 7 | 99.70 | 99.94 | 99.64 |
| *Lt foot* | 6617 | 14 | 171 | 16 | 99.79 | 97.48 | 97.29 | 5915 | 11 | 605 | 70 | 99.81 | 90.72 | 90.67 |
| **Overall** |  |  |  |  | 99.56 | 89.24 | 88.96 |  |  |  |  | 99.51 | 88.11 | 87.78 |

**Table S11.** Fold_3, trained on data from infants # 1, 2, 3, 5, 6;

|  | *Test results on video #2 of baby 4* |  |  |  |  |  |  | *Test results on video #1 of baby 4* |  |  |  |  |  |  |
| --- | --- | --- | --- | --- | --- | --- | --- | --- | --- | --- | --- | --- | --- | --- |
|  | *TP* | *FP* | *FN* | *TN* | *Selectivity* | *Sensitivity* | *Accuracy* | *TP* | *FP* | *FN* | *TN* | *Selectivity* | *Sensitivity* | *Accuracy* |
| *Rt shoulder* | 7455 | 1 | 0 | 0 | 99.99 | 100.00 | 99.99 | 7145 | 5 | 4 | 0 | 99.93 | 99.94 | 99.87 |
| *Rt elbow* | 7429 | 26 | 1 | 0 | 99.65 | 99.99 | 99.64 | 7146 | 7 | 1 | 0 | 99.90 | 99.99 | 99.89 |
| *Rt hand* | 7161 | 37 | 230 | 28 | 99.49 | 96.89 | 96.42 | 6982 | 30 | 122 | 20 | 99.57 | 98.28 | 97.88 |
| *Rt hip* | 7199 | 6 | 190 | 61 | 99.92 | 97.43 | 97.37 | 6217 | 5 | 842 | 90 | 99.92 | 88.07 | 88.16 |
| *Rt knee* | 7044 | 34 | 353 | 25 | 99.52 | 95.23 | 94.81 | 5907 | 3 | 1157 | 87 | 99.95 | 83.62 | 83.79 |
| *Rt foot* | 5997 | 21 | 1381 | 57 | 99.65 | 81.28 | 81.20 | 6496 | 22 | 591 | 45 | 99.66 | 91.66 | 91.43 |
| *Lt shoulder* | 7452 | 4 | 0 | 0 | 99.95 | 100.00 | 99.95 | 7151 | 3 | 0 | 0 | 99.96 | 100.00 | 99.96 |
| *Lt elbow* | 7389 | 16 | 38 | 13 | 99.78 | 99.49 | 99.28 | 7120 | 30 | 1 | 3 | 99.58 | 99.99 | 99.57 |
| *Lt hand* | 6473 | 12 | 906 | 65 | 99.81 | 87.72 | 87.69 | 5292 | 17 | 1781 | 64 | 99.68 | 74.82 | 74.87 |
| *Lt hip* | 7234 | 7 | 146 | 69 | 99.90 | 98.02 | 97.95 | 6940 | 15 | 150 | 49 | 99.78 | 97.88 | 97.69 |
| *Lt knee* | 4658 | 6 | 2751 | 41 | 99.87 | 62.87 | 63.02 | 5606 | 12 | 1466 | 70 | 99.79 | 79.27 | 79.34 |
| *Lt foot* | 7128 | 73 | 237 | 18 | 98.99 | 96.78 | 95.84 | 6592 | 63 | 458 | 41 | 99.05 | 93.50 | 92.72 |
| *Overall* |  |  |  |  | 99.71 | 92.97 | 92.76 |  |  |  |  | 99.73 | 92.25 | 92.10 |

**Table S11.** Fold_4, trained on data from infants # 1, 2, 4, 4, 6;

|  | **Test results on video #2 of baby 3** |  |  |  |  |  |  | **Test results on video #1 of baby 3** |  |  |  |  |  |  |
| --- | --- | --- | --- | --- | --- | --- | --- | --- | --- | --- | --- | --- | --- | --- |
|  | *TP* | *FP* | *FN* | *TN* | *Selectivity* | *Sensitivity* | *Accuracy* | *TP* | *FP* | *FN* | *TN* | *Selectivity* | *Sensitivity* | *Accuracy* |
| *Rt shoulder* | 6277 | 3 | 47 | 25 | 99.95 | 99.26 | 99.21 | 6113 | 1 | 23 | 18 | 99.98 | 99.63 | 99.61 |
| *Rt elbow* | 6073 | 12 | 199 | 68 | 99.80 | 96.83 | 96.68 | 5348 | 1 | 724 | 82 | 99.98 | 88.08 | 88.22 |
| *Rt hand* | 6051 | 10 | 223 | 68 | 99.84 | 96.45 | 96.33 | 6042 | 10 | 73 | 30 | 99.83 | 98.81 | 98.65 |
| *Rt hip* | 5560 | 0 | 687 | 105 | 100.00 | 89.00 | 89.18 | 6106 | 5 | 14 | 30 | 99.92 | 99.77 | 99.69 |
| *Rt knee* | 6266 | 38 | 23 | 25 | 99.40 | 99.63 | 99.04 | 6131 | 4 | 10 | 10 | 99.93 | 99.84 | 99.77 |
| *Rt foot* | 6303 | 11 | 27 | 11 | 99.83 | 99.57 | 99.40 | 6149 | 2 | 3 | 1 | 99.97 | 99.95 | 99.92 |
| *Lt shoulder* | 6348 | 1 | 3 | 0 | 99.98 | 99.95 | 99.94 | 6125 | 3 | 15 | 12 | 99.95 | 99.76 | 99.71 |
| *Lt elbow* | 6307 | 8 | 17 | 20 | 99.87 | 99.73 | 99.61 | 6014 | 31 | 73 | 37 | 99.49 | 98.80 | 98.31 |
| *Lt hand* | 6335 | 8 | 4 | 5 | 99.87 | 99.94 | 99.81 | 6127 | 8 | 12 | 8 | 99.87 | 99.80 | 99.68 |
| *Lt hip* | 6248 | 5 | 58 | 41 | 99.92 | 99.08 | 99.01 | 5934 | 9 | 132 | 80 | 99.85 | 97.82 | 97.71 |
| *Lt knee* | 5891 | 17 | 356 | 88 | 99.71 | 94.30 | 94.13 | 5112 | 8 | 978 | 57 | 99.84 | 83.94 | 83.98 |
| *Lt foot* | 6231 | 5 | 96 | 20 | 99.92 | 98.48 | 98.41 | 6107 | 1 | 35 | 12 | 99.98 | 99.43 | 99.42 |
| **Overall** |  |  |  |  | 99.84 | 97.69 | 97.56 |  |  |  |  | 99.88 | 97.14 | 97.06 |

**Table S12.** Fold_5, trained on data from infants # 1, 3, 4, 5, 6;

|  | *Test results on video #2 of baby 2* |  |  |  |  |  |  | *Test results on video #1 of baby 2* |  |  |  |  |  |  |
| --- | --- | --- | --- | --- | --- | --- | --- | --- | --- | --- | --- | --- | --- | --- |
|  | *TP* | *FP* | *FN* | *TN* | *Selectivity* | *Sensitivity* | *Accuracy* | *TP* | *FP* | *FN* | *TN* | *Selectivity* | *Sensitivity* | *Accuracy* |
| *Rt shoulder* | 5942 | 2 | 8 | 1 | 99.97 | 99.87 | 99.83 | 5672 | 1 | 0 | 2 | 99.98 | 100.00 | 99.98 |
| *Rt elbow* | 5875 | 6 | 47 | 25 | 99.90 | 99.21 | 99.11 | 5613 | 4 | 44 | 14 | 99.93 | 99.22 | 99.15 |
| *Rt hand* | 5824 | 44 | 77 | 8 | 99.25 | 98.70 | 97.97 | 5574 | 91 | 8 | 2 | 98.39 | 99.86 | 98.26 |
| *Rt hip* | 5940 | 0 | 8 | 5 | 100.00 | 99.87 | 99.87 | 5547 | 2 | 87 | 39 | 99.96 | 98.46 | 98.43 |
| *Rt knee* | 5941 | 11 | 1 | 0 | 99.82 | 99.98 | 99.80 | 5667 | 8 | 0 | 0 | 99.86 | 100.00 | 99.86 |
| *Rt foot* | 5855 | 3 | 83 | 12 | 99.95 | 98.60 | 98.56 | 5659 | 15 | 0 | 1 | 99.74 | 100.00 | 99.74 |
| *Lt shoulder* | 5918 | 12 | 18 | 5 | 99.80 | 99.70 | 99.50 | 5609 | 4 | 41 | 21 | 99.93 | 99.27 | 99.21 |
| *Lt elbow* | 5315 | 1 | 557 | 80 | 99.98 | 90.51 | 90.63 | 5590 | 0 | 58 | 27 | 100.00 | 98.97 | 98.98 |
| *Lt hand* | 5767 | 17 | 139 | 30 | 99.71 | 97.65 | 97.38 | 4831 | 3 | 794 | 47 | 99.94 | 85.88 | 85.96 |
| *Lt hip* | 5944 | 5 | 3 | 1 | 99.92 | 99.95 | 99.87 | 2715 | 0 | 2915 | 45 | 100.00 | 48.22 | 48.63 |
| *Lt knee* | 5930 | 23 | 0 | 0 | 99.61 | 100.00 | 99.61 | 5652 | 23 | 0 | 0 | 99.59 | 100.00 | 99.59 |
| *Lt foot* | 5931 | 4 | 12 | 6 | 99.93 | 99.80 | 99.73 | 5624 | 6 | 34 | 11 | 99.89 | 99.40 | 99.30 |
| *Overall* |  |  |  |  | 99.82 | 98.65 | 98.49 |  |  |  |  | 99.77 | 94.11 | 93.92 |

**Table S13.** Fold_6, trained on data from infants # 2, 3, 4, 5, 6;

|  | *Test results on video #2 of baby 1* |  |  |  |  |  |  | *Test results on video #1 of baby 1* |  |  |  |  |  |  |
| --- | --- | --- | --- | --- | --- | --- | --- | --- | --- | --- | --- | --- | --- | --- |
|  | *TP* | *FP* | *FN* | *TN* | *Selectivity* | *Sensitivity* | *Accuracy* | *TP* | *FP* | *FN* | *TN* | *Selectivity* | *Sensitivity* | *Accuracy* |
| *Rt shoulder* | 1213 | 3 | 697 | 21 | 99.75 | 63.51 | 63.81 | 1784 | 4 | 58 | 6 | 99.78 | 96.85 | 96.65 |
| *Rt elbow* | 633 | 0 | 1292 | 9 | 100.00 | 32.88 | 33.20 | 1836 | 1 | 12 | 3 | 99.95 | 99.35 | 99.30 |
| *Rt hand* | 1012 | 3 | 914 | 5 | 99.70 | 52.54 | 52.59 | 697 | 0 | 1152 | 3 | 100.00 | 37.70 | 37.80 |
| *Rt hip* | 1325 | 2 | 586 | 21 | 99.85 | 69.34 | 69.60 | 1400 | 2 | 438 | 12 | 99.86 | 76.17 | 76.24 |
| *Rt knee* | 1920 | 13 | 1 | 0 | 99.33 | 99.95 | 99.28 | 1788 | 12 | 45 | 7 | 99.33 | 97.55 | 96.92 |
| *Rt foot* | 1835 | 11 | 84 | 4 | 99.40 | 95.62 | 95.09 | 1793 | 3 | 49 | 7 | 99.83 | 97.34 | 97.19 |
| *Lt shoulder* | 1810 | 3 | 98 | 23 | 99.83 | 94.86 | 94.78 | 1784 | 1 | 62 | 5 | 99.94 | 96.64 | 96.60 |
| *Lt elbow* | 1425 | 2 | 490 | 17 | 99.86 | 74.41 | 74.56 | 1802 | 12 | 34 | 4 | 99.34 | 98.15 | 97.52 |
| *Lt hand* | 864 | 3 | 1059 | 8 | 99.65 | 44.93 | 45.09 | 1010 | 3 | 831 | 8 | 99.70 | 54.86 | 54.97 |
| *Lt hip* | 1045 | 0 | 880 | 9 | 100.00 | 54.29 | 54.50 | 1335 | 0 | 500 | 17 | 100.00 | 72.75 | 73.00 |
| *Lt knee* | 1849 | 16 | 66 | 3 | 99.14 | 96.55 | 95.76 | 1833 | 14 | 2 | 3 | 99.24 | 99.89 | 99.14 |
| *Lt foot* | 945 | 0 | 982 | 7 | 100.00 | 49.04 | 49.22 | 1425 | 1 | 416 | 10 | 99.93 | 77.40 | 77.48 |
| *Overall* |  |  |  |  | 99.71 | 68.99 | 68.95 |  |  |  |  | 99.74 | 83.72 | 83.57 |

**Figure S2.**
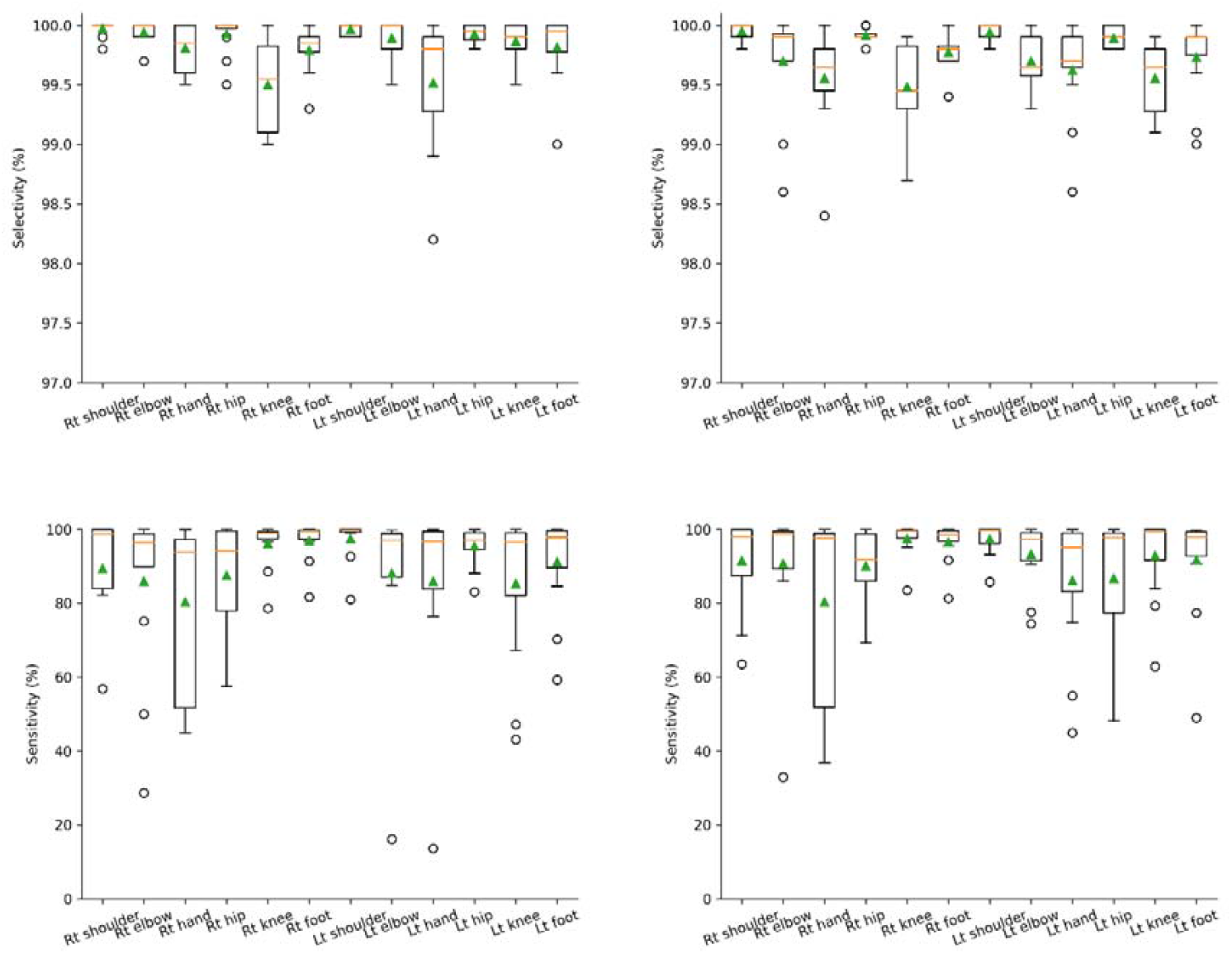
Precision and sensitivity measures for each anatomical location using the resnet (A and C) and the efficientnet (B and D), respectively. Circles indicate outliers. Green triangles represent mean values.

**Figure S3.**
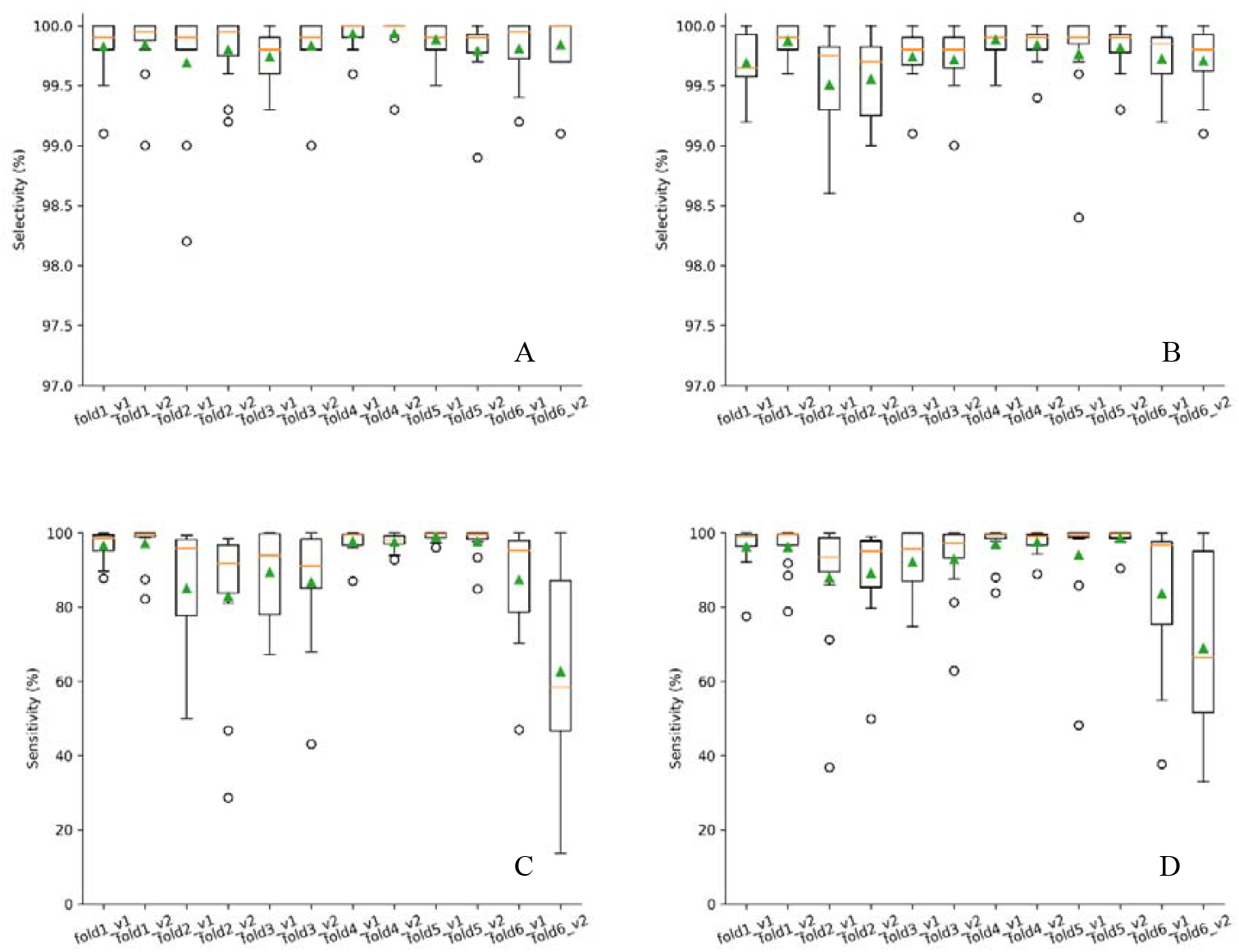
Precision and sensitivity measures for each validation set (unseen videos) using the resnet (A and C) and the efficientnet (B and D), respectively. Circles indicate outliers. Green triangles represent mean values.

**Figure S4.**
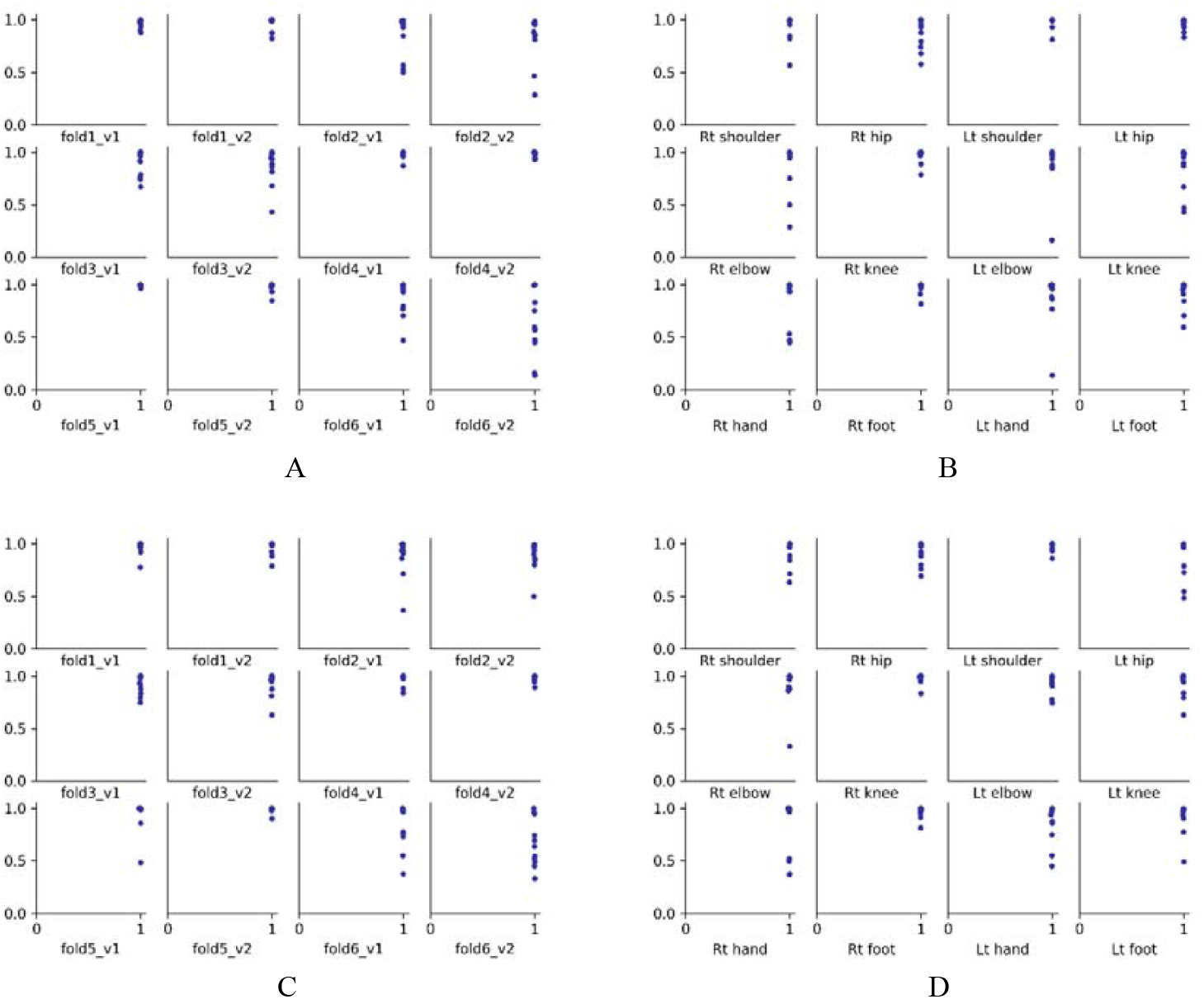
Scattered precision-recall plots across the 12 anatomical locations for all tested videos using Resnet (A) and Efficient_net (C), and across the novel videos in the validation sets for the 12 body parts from the unseen infants (two videos per fold), using Resnet (B) and Efficient_net (D).

**Figure S5.**
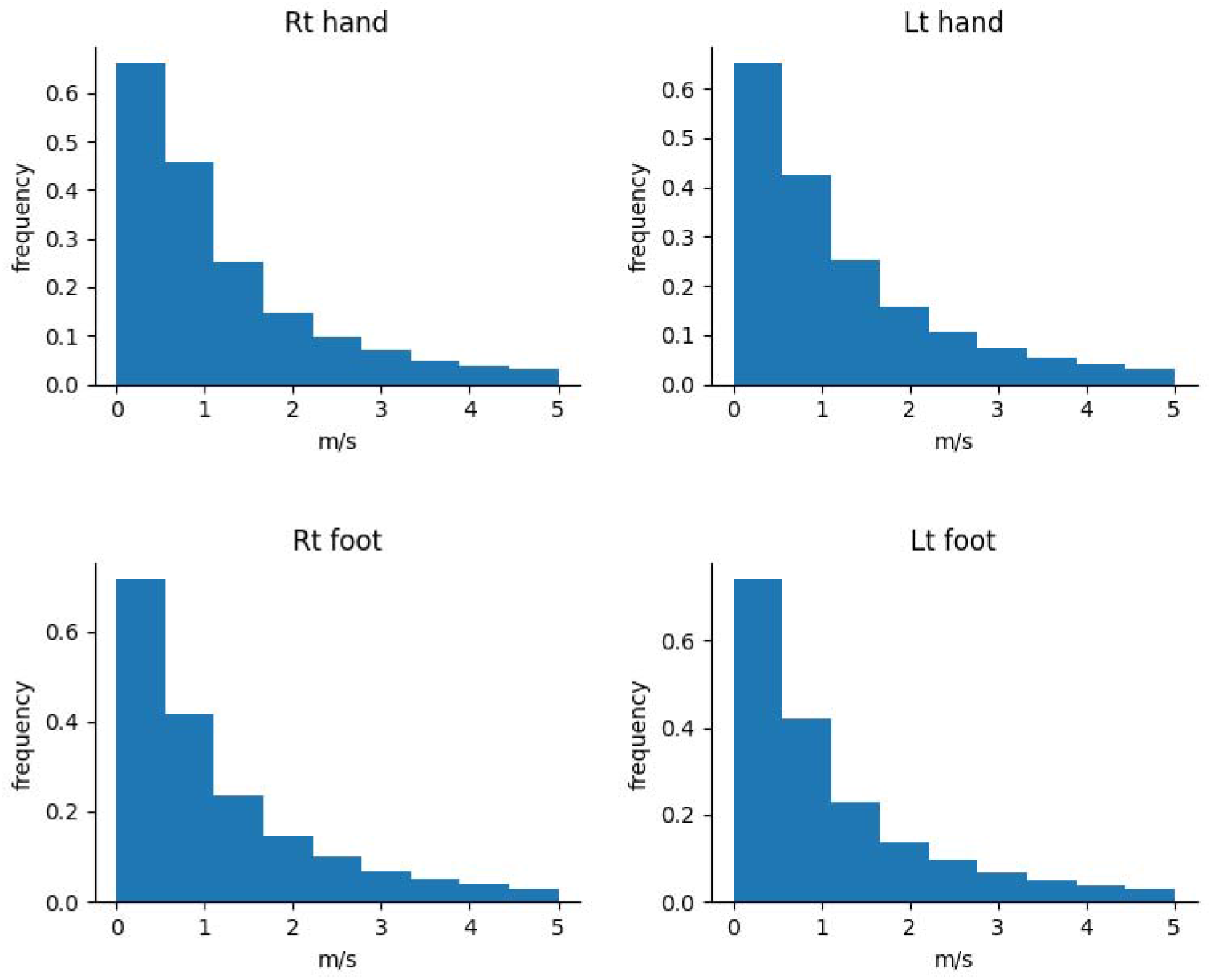
Normalized histograms of the hands and feets’ velocity across all 12 videos from all participants, calculated from DLC output with 1-frame time step.

